# FiberLM: A Transformer-Based Model for Mouse Brain Diffusion MRI Tractography Guided by Viral Tracer Data

**DOI:** 10.64898/2026.05.06.723316

**Authors:** Ray Wen, Jiangyang Zhang, Zifei Liang

## Abstract

Diffusion MRI (dMRI) tractography provides a non-invasive method for mapping whole-brain structural connectivity. However, its application is limited by substantial false-positive and false-negative connections. While deep learning based methods have shown promise in improving tractography, most rely on training data derived from conventional dMRI tractography, therefore inheriting the same limitations. Here, we introduce FiberLM, an attention-based Transformer model for mouse brain tractography. The model was trained using a whole-brain streamline dataset based on viral tracer data from the Allen Mouse Brain Connectivity Atlas (AMBCA), allowing the model to learn the properties of both local and long-range axonal trajectories through self-attention. FiberLM was applied to predict anatomically plausible axonal trajectories from ex vivo high-resolution mouse brain dMRI data. Quantitative evaluations demonstrated that FiberLM significantly reduced false-positive and false-negative connections, improved spatial agreement with tracer-defined pathways, and generated whole-brain connectomes that more closely approximated AMBCA results compared to conventional tractography. These findings suggest FiberLM as a potential tool for accurate reconstruction of mouse brain structural connectomics.

## Introduction

Diffusion magnetic resonance imaging (dMRI) tractography provides the only non-invasive approach to reconstructing the brain’s axonal pathways at a whole-brain scale (Mori and Van Zijl, 2002; Jeurissen et al., 2019). By mapping the “wiring diagram” of the brain, tractography offers unprecedented insights into the structural connectivity that underlies cognition, development, and disease (Jbabdi and Johansen-Berg, 2011; Sporns et al., 2005). Tractography typically begins by estimating local axonal fiber orientations from dMRI signals and then propagates streamlines following the fiber orientations. These tractography streamlines represent potential axonal pathways in the brain. Early tractography methods relied on diffusion tensor imaging (Basser et al., 1994) for fiber orientation information, and techniques such as Q-ball (Tuch et al., 2002) and high angular resolution diffusion imaging (HARDI)(Frank, 2001) were introduced later to better handle complex fiber configurations such as crossing fibers (Descoteaux et al., 2007; Tournier et al., 2007). Three major streamline propagation methods have been developed. Deterministic tracking follows the primary fiber orientation at each voxel, readily capturing large axonal pathways but is vulnerable to error accumulation and crossing fibers (Mori et al., 1999; Mori and Van Zijl, 2002). Probabilistic tracking samples from local fiber orientation distributions to generate multiple streamlines, improving sensitivity to small or crossing axonal bundles, though at the risk of generating spurious connections (Behrens et al., 2003; Parker et al., 2003). Global tractography reconstructs streamlines by minimizing a global energy function, improving anatomical coherence but at high computational cost and depends on the choice of energy function (Fillard et al., 2011; Mangin et al., 2013; Reisert et al., 2011). Yet, despite methodological progress, tractography remains prone to substantial false-positive and false-negative connections (Maier-Hein et al., 2017; Rheault et al., 2020; Schilling et al., 2019a), which hinders the applications of tractography in neuroscience research.

To address these limitations, anatomically and microstructurally constrained methods were developed to reduce spurious reconstructions. Anatomically constrained tractography incorporated anatomical priors to restrict streamline propagation to plausible pathways (Huang et al., 2004; Smith et al., 2012). In addition, parcellation- or bundle-specific approaches regularized streamlines according to known axonal trajectories (Girard et al., 2014; Theaud et al., 2020; Wasserthal et al., 2018). Microstructure-informed techniques integrated biophysical models to refine fiber orientation estimates (Daducci et al., 2016). Streamline clustering and filtering methods, such as FIbEr Segmentation in Tractography using Autoencoders (FIESTA), grouped anatomically consistent fibers to suppress false positives (Dumais et al., 2023). However, these methods remain constrained by their reliance on dMRI-derived fiber orientation distribution functions (fODFs) and existing streamline propagation algorithms, which do not fully address the risk of false negatives or inaccurate axonal pathway reconstruction.

The emergence of machine learning and deep learning methods introduced new opportunities to reduce false positive and negative connections in dMRI tractography. For example, residual block deep neural network (ResDNN) has been used to estimate fiber orientation from dMRI signals (Nath et al., 2019), and recurrent neural networks (RNNs) have been used to improve streamline propagation by leveraging potential sequential dependencies (Benou and Riklin-Raviv, 2018; Poulin et al., 2019). Other approaches include framing tracking as a classification (Legarreta et al., 2021; Neher et al., 2015), or sequential decision-making process with reinforced learning (Théberge et al., 2021). More recently, attention-based architectures, particularly the Transformer architecture (Vaswani et al., 2017), have been introduced to capture long-range dependencies and integrate global anatomical context across entire streamline sequences (Waizman et al., 2025; Yang et al., 2025). These studies demonstrated the potential of machine learning and deep learning based tractography, but training deep networks solely on diffusion-derived fODFs or tractograms risks inheriting the same biases that underlie conventional methods (Aydogan et al., 2018; Grisot et al., 2021; Schilling et al., 2019b; Seehaus et al., 2013), motivating the use of independent ground-truth data for model supervision.

Chemical and viral tracer data imaged with light microscopy have emerged as a promising source of such ground truth. Several reports have shown that using neural networks trained on tracer data can yield more accurate fODF estimates than conventional methods (Schilling et al., 2018; Nath et al., 2019; Liang et al., 2023; Zhu et al., 2025). In our previous work, we combined streamline data from thousands of individual tracer experiments from the Allen Mouse Brain Connectivity Atlas (AMBCA) (Kuan et al., 2015; Oh et al., 2014) to construct a comprehensive whole-brain representation of mouse forebrain connectivity at the macroscopic and mesoscopic levels (Liang et al., 2023). Here, we leveraged this resource to introduce FiberLM, a streamline propagation model for mouse brain tractography that integrates the transformer architecture with tracer-informed supervision. By encoding fODFs as sequential tokens, we aimed to capture both local and long-range structural dependencies through self-attention, enabling anatomically grounded whole-brain tractography.

## 2. Method

### 2.1 AMBCA streamlines dataset

A mesoscale streamline dataset was constructed from AMBCA (Kuan et al., 2015), which provides axonal projections reconstructed from anterograde viral tracer experiments across thousands of mouse brains. As described in (Liang et al., 2023), streamlines from all experiments were aggregated in the Allen Reference Atlas (ARA) space. To address hemispheric sampling bias introduced by predominantly unilateral tracer injections, the aggregated streamlines were mirrored across the midline, yielding a symmetric whole-brain tractogram comprising 4 million streamlines. The dataset can be downloaded at https://github.com/liangzifei/MR-TOD-net.

### 2.2 *Ex vivo* dMRI

Mouse brain dMRI data were acquired from a separate cohort of adult mice. All animal experiments have been approved by the Institute Animal Care and Use Committee at New York University. To match the AMBCA data, adult C57BL/6 mice (7-8 weeks old, n = 10, 5M/5F, Charles River, Wilmington, MA, USA) were perfusion fixed with 4% paraformaldehyde (PFA) in PBS. The samples were preserved in 4% PFA for 24 h before transferring to PBS. Ex vivo MRI of mouse brain specimens was performed on a horizontal 7 Tesla MR scanner (Bruker Biospin, Billerica, MA, USA) with a triple-axis gradient system. Images were acquired using a quadrature volume excitation coil (72 mm inner diameter) and a receive-only 4-channel phased array cryogenic coil. The specimens were imaged with the skull intact and placed in a syringe filled with Fomblin (perfluorinated polyether, Solvay Specialty Polymers USA, LLC, Alpharetta, GA, USA) to prevent tissue dehydration (Arefin et al., 2021). Three-dimensional dMRI data were acquired using a modified 3D diffusion-weighted gradient- and spin-echo (DW-GRASE) sequence (Aggarwal et al., 2010; Wu et al., 2013) with the following parameters: echo time (TE)/repetition time (TR) = 30/400 ms; two signal averages; field of view (FOV) = 12.8 mm × 10 mm × 18 mm, resolution = 0.1 mm × 0.1 mm × 0.1 mm; two non diffusion weighted images (b0s); 60 diffusion weighted images (DWIs) with a diffusion weighting (b) of 5,000 s/mm^2^.

### 2.3 dMRI data analysis

From the ex vivo dMRI data, diffusion tensors were estimated at each voxel using log-linear fitting implemented in MRtrix3 (http://www.mrtrix.org)(Tournier et al., 2012), and maps of mean diffusivity (MD) and fractional anisotropy (FA) were generated. Fiber orientation distributions (FODs) were estimated using constrained spherical deconvolution (CSD) with a single response function derived from representative white matter voxels (Tournier et al., 2007, 2004). For consistent cross-subject comparisons, all tractography experiments were conducted in the ARA space. Mappings between individual mouse brain images and the Allen Reference Atlas (ARA) space were computed using Large Deformation Diffeomorphic Metric Mapping (LDDMM) (Beg et al., 2005) as described in (Arefin et al., 2021). The LDDMM-generated deformation fields were converted into MRtrix-compatible warp format, and FODs were spatially transformed into ARA space using the *mrtransform* command together with the reorientation of FOD (Raffelt et al., 2011), which adjust the FODs based on the deformation field.

### 2.4 Whole Brain tractography and connectivity map construction

Whole-brain probabilistic tractography based on the re-oriented FODs was performed using MRtrix3 (Tournier et al., 2012) with the second-order integration over fiber orientation distributions (iFOD2) algorithm. Streamlines were seeded randomly within the whole-brain mask and propagated using the following parameters: step size = 0.025 mm, maximum turning angle = 45°, minimum streamline length = 3 mm, and FOD amplitude cutoff = 0.05. Whole-brain tractograms were generated with streamline counts ranging from 100,000 (for qualitative visualization and methodological comparison) to 1 million streamlines for structural connectome construction and quantitative analyses.

Structural connectivity matrices were constructed based on the 184 regions selected from the ARA atlas. Each streamline was assigned to a pair of brain regions according to its terminal endpoints. Streamline counts between region pairs were aggregated using tck2connectome to produce symmetric, zero-diagonal adjacency matrices representing the mouse brain structural connectome. Streamline counts were used to measure inter-regional structural connectivity. For nodal-level analyses, streamlines passing through a given region of interest (ROI) were isolated using *tckedit* with the -include option, enabling assessment of node-specific connectivity patterns and tractographic topology.

### 2.5 The architecture of FiberLM, an attention-Based Transformer

To capture both local FODs and long-range contextual dependencies along streamlines, we implemented an attention-based Transformer decoder network (FiberLM) in PyTorch. Each streamline was represented as an ordered sequence of FODs,

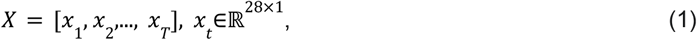

where each *x*_*t*_ corresponds to the 28-dimensional spherical harmonic (SH) representation of the FOD at the streamline position *t*. These feature sequences were used as sequential input tokens for the FiberLM, which was designed to predict the probability distribution of the next propagation direction *d*_*t*_ on a 724-point unit sphere plus one termination class. The network architecture follows a decoder-only Transformer design (**Fig. 1A**), consisting of two main components:

1. **Embedding and streamline positional encoding:** Each 28-dimensional FOD coefficient vector is first linearly projected into a 1024-dimensional embedding space (*d*_*model*_), followed by layer normalization. A sinusoidal positional encoding, as used in language model (Vaswani et al., 2017), is then added to each FOD vertex embedding, injecting sequential order information and enabling the model to distinguish similar local FOD features occurring at different steps along the streamline. This combined representation is subsequently normalized through layer normalization. The resulting streamline position-aware representation is denoted 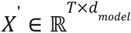, where *T* = 128 is the fixed sequence length to which all streamlines are zero-padded or truncated.
2. **Context integration block:** *X*′ is then processed through twelve stacked decoder layers. Each layer consists of a masked multi-head self-attention sublayer (16 heads, *d*_*k*_ = *d*_*v*_ = 64, dropout = 0.1) followed by a two-layer feed-forward sublayer (dropout = 0.1), with pre-layer normalization and residual connections applied around each sublayer. The causal attention mask restricts each position to attend only to itself and preceding positions, preserving autoregressive consistency between training and inference.

**Figure 1.**
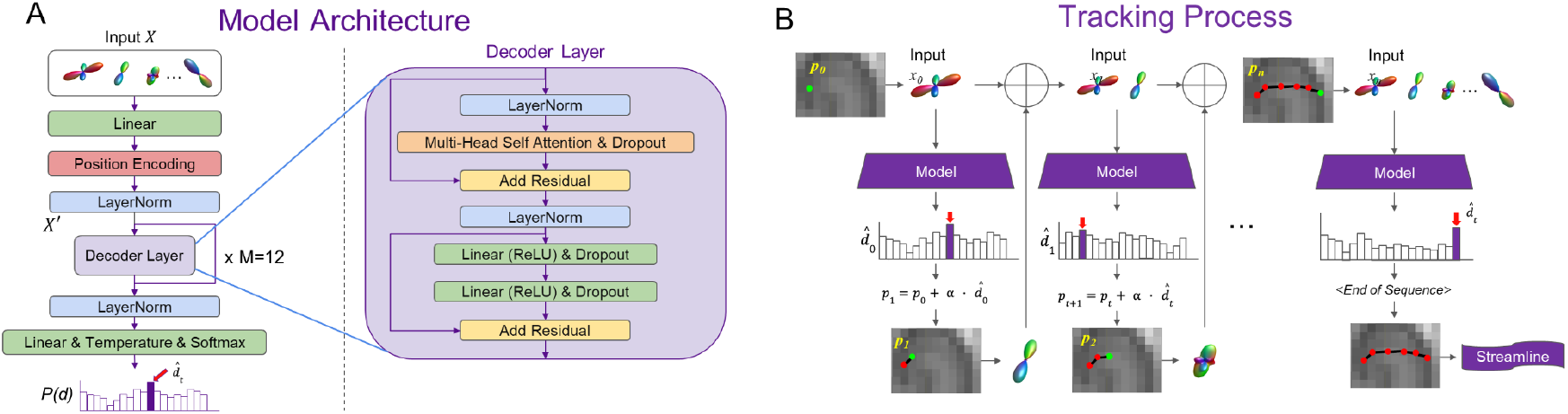
Deep network architecture and tracking process. (**A**) Transformer-based decoder architecture for streamline propagation, with embedded inputs processed by stacked self-attention and feed-forward layers to predict direction probabilities. (**B**) Autoregressive tracking procedure in which predicted directions are sequentially sampled and integrated to form a complete streamline.

Finally, after a concluding layer normalization, a linear projection converts the decoder output into unnormalized logits over discrete orientation directions (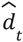, **Fig. 1A)**. During tracking, these logits are scaled by a temperature factor of 0.5 and normalized via softmax (Brown et al., 2020) to yield the probability distribution *P(d)*(**Fig. 1B**).

The attention mechanism is defined as

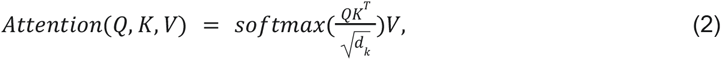

Where the query (*Q*), key (*K*), and value (*V*) are obtained by projecting *X*′ with:

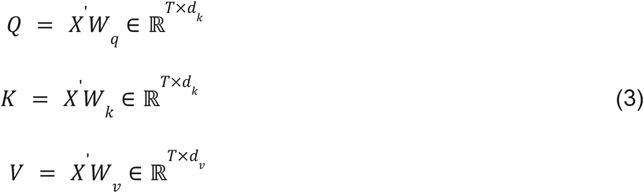

where 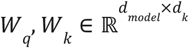 and 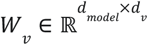 are learned projection matrices that map the input into the attention subspace. In the multi-head formulation (**Fig. 1A**, Decoder Layer), this scaled dot-product attention is performed independently across all 16 parallel heads. The resulting *T* × *d*_*v*_ outputs are concatenated along the feature dimension and linearly projected back to the original *d*_*model*_ dimension via a learned output weight matrix. This formulation allows the network to capture long-range dependencies across all preceding streamline steps while retaining sensitivity to local FOD orientation at each position. During inference, the trained Transformer receives 1 million seed points within the whole-brain mask. At each propagation step, the model receives the accumulated FOD sequence and outputs *P(d)*. The next-step orientation 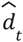 is then selected as the direction with the highest probability in *P(d)*. Streamlines are advanced with a fixed step size of 0.05 mm, and tracking terminates upon entering non-white-matter tissue, exceeding a 60° curvature threshold, falling outside the valid length range of 1 to 10 mm in length, or triggering entropy-based tracking termination (Benou and Riklin-Raviv, 2018), which halts propagation when the output distribution indicates high directional uncertainty. Because generated streamlines are routinely discarded for violating these constraints, the seed sampling process was repeated until approximately 1 million valid streamlines were generated.

### 2.6 Model training and implementation details (Hyperparameters)

Training samples were generated from the AMBCA streamline dataset by extracting a sequence of FOD features from the orientation-adjusted FOD data normalized to the ARA space for each streamline and corresponding target directions. Data from eight subjects were used, with four for training and four held out for independent testing; 3 million streamlines were randomly sampled from the training subjects and pooled to form the training set. To enable both forward and backward streamline generation, each AMBCA streamline was augmented by direction reversal. The FOD peak amplitudes were normalized across subjects, ensuring consistency in feature scaling and directional representation across the training dataset. The model was optimized with the Adam optimizer (weight decay = 0.001, β_1_ = 0.9, β_2_ = 0.999, ε = 1 × 10^-7^, clipnorm = 1.0) using a batch size of 128. The model was trained for a total of 6 epochs. During the first 5 epochs, the learning rate followed a cosine decay schedule, beginning with a linear warmup over the first 6% of steps (31,772 steps) to reach a peak learning rate of 1. 5 × 10^−4^. The learning rate then decayed over 497,773 steps to a minimum of 1. 5 × 10^−5^. During the 6th and final epoch, the learning rate was held constant at this minimum value of 1. 5 × 10^−5^.

A categorical cross-entropy loss was applied to optimize the fiber-class predictions. The loss was computed exclusively over the valid (non-padded) vertices to prevent artificial attenuation.

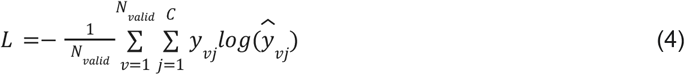

Where *C* is the total number of discrete orientation classes, *N*_*valid*_ is the total number of valid vertices across the training batch. For a given vertex *v, y*_*vj*_ ∈ {0, 1} is the ground-truth binary indicator for class *j*, and 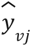 is the corresponding predicted probability from *P(d)*.

All experiments were conducted on an NVIDIA A100 GPU (80GB VRAM) with CUDA 11.8. The final trained network demonstrated stable convergence and generalization across subjects, supporting robust whole-brain tractography in mouse datasets.

### 2.7 Attention weighting and streamline curvature

For a streamline with *T* vertices, each decoder layer *l* and head *h* produces an attention weight matrix *A*^(*l,h*)^ ∈ ℝ^*T*×*T*^ computed via scaled dot-product attention with row-wise softmax normalization (Javed et al., 2025; Lee et al., 2025). First, the attention weights were aggregated across all heads and layers by simple averaging to create a single aggregated attention matrix:

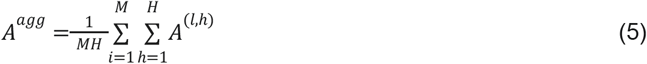

where *M* = 12 and *H*=16 denote the number of decoder layers and attention heads, respectively. To ensure a consistent mean attention weight of 1 across all rows, *A*^*agg*^ was scaled such that each valid cell in row *i* is multiplied by *i*. The scaled attention matrix 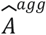 is defined as:

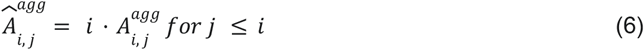

where *j* ≤ *i* represents the valid, unmasked tokens in the causal sequence. A scalar importance score for each streamline segment *j* was computed as the masked mean of the incoming attention. To prevent artificial dilution from causal masking, the sum of incoming attention was divided only by the number of valid (non-zero) elements in column *j*, denoted as *N*_*valid*_ (*j*):

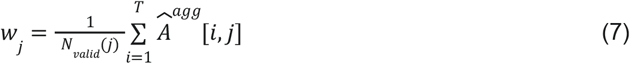

Each segment weight *w*_*j*_ was mapped to template space, and voxelwise attention was obtained by averaging weights of all streamline segments within each voxel *v*:

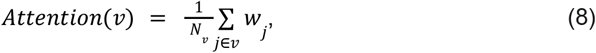

where *N*_*v*_ is the number of streamline segments within voxel *v*. The resulting map was normalized to the [0,1] range across the brain.

### 2.8 Data and statistical analysis

Connectivity matrices generated using the t*ck2connectome* command in MRtrix were converted to binary matrices using a threshold of 0.001 times the maximal amplitude (Arefin et al., 2023). Four global metrics were calculated using the Brain Connectivity Toolbox (BCT) (Rubinov and Sporns, 2010): global efficiency, clustering coefficient, modularity, and characteristic path length. Local geometric complexity of streamlines was quantified using the curvature contrast implemented in MRtrix3 (*tckmap*-contrast curvature). In this framework, streamline curvature is computed directly from the discrete geometry of each streamline during tract density image (TDI) generation. During voxel-wise mapping, curvature values from all streamline samples traversing a given voxel are accumulated and averaged. The resulting curvature map therefore reflects the mean local bending of fiber trajectories within each voxel, with higher values indicating greater geometric complexity.

To compare conventional and FiberLM generated tractography results from the same subjects, we used paired t-tests with Benjamini–Hochberg false discovery rate (FDR) corrections (q < 0.05). The Dice similarity coefficient was used to quantify the spatial overlap between tractography-derived bundles and the AMBCA streamlines between a pair of regions, using binarized voxel maps of streamlines (Arefin et al., 2023). The false positive rate and false negative rate were computed as the proportions of connections unique to tractography or the reference relative to the total reference connections. Average false positive and negative rates across subjects were also compared for each region to map whole-brain distributions of false connections.

## 3. Results

### 3.1. Whole-brain tractograms generated by FiberLM recapitulated the spatial connectivity patterns of the AMBCA streamline dataset

**Fig. 2A** compares whole mouse brain tractograms generated by FiberLM and conventional probabilistic tractography using MRtrix with 0.1 million and 1 million total streamlines. The dMRI data were not used in training. At 0.1 million streamlines, FiberLM-generated tractograms showed greater cortical and hippocampal coverage than conventional probabilistic tractography but had sparse streamlines in the cerebellum (indicated by the white arrow in **Fig. 2A**). As the streamline count increased to 1 million, FiberLM-generated tractograms exhibited improved cerebellar coverage, while conventional probabilistic tractography showed increased cortical and hippocampal streamline density. Major white matter pathways in FiberLM-generated tractograms were consistent with probabilistic tractography results and corresponding tracts in the AMBCA streamline dataset. Three representative tracts, the corpus callosum, cerebral peduncle, and anterior commissure, were isolated from whole-brain tractograms using identical regions of interest and showed comparable trajectories across methods (**Fig. 2B**). Notably, conventional probabilistic tractography yielded a predominantly radial orientation pattern in the cortex and hippocampus (indicated by the red and yellow arrow in **Fig. 2A**), whereas FiberLM recapitulated the subtle orientation variations observed in the AMBCA streamline dataset.

**Figure 2:**
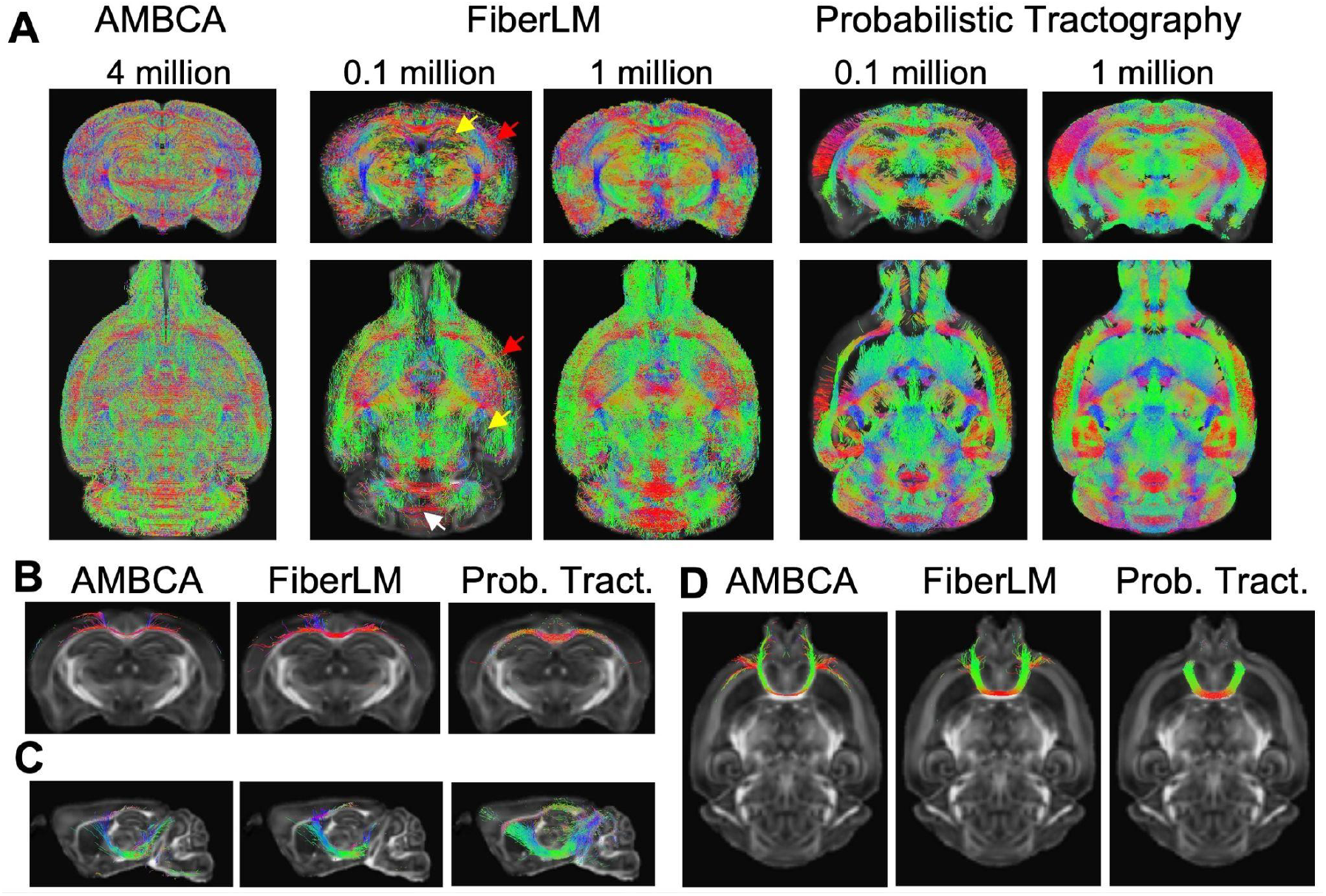
Comparison of FiberLM and probabilistic tractography results against AMBCA streamlines. (**A**) Whole mouse brain tractograms reconstructed using FiberLM and probabilistic tractography at different total streamline counts were compared with the AMBCA streamlines. The streamlines are overlaid on the FA images. The images on the top are coronal views of the brain, and the images on the bottom are horizontal views of the brain. The red, yellow, and white arrows indicate the locations of the cortex, hippocampus, and cerebellum. **(B-D)** The corpus callosum, cerebral peduncle, and anterior commissure selected from the above tractograms and the corresponding AMBCA streamlines. Streamlines are color-coded by local orientations (red: left–right; green: anterior–posterior; blue: superior–inferior).

### 3.2. Whole-brain connectivity reconstructed using FiberLM better preserved the network properties than conventional probabilistic tractography

**Fig. 3** compares connectivity matrices derived from the AMBCA streamline dataset with those generated by FiberLM and conventional probabilistic tractography. The nodes in the matrices are organized into nine major brain regions: cortical subregions, cortical subplate, hippocampus, pallidum, striatum, thalamus, hypothalamus, midbrain, and hindbrain. Overall, both FiberLM and probabilistic tractography recapitulated the large-scale modular organization observed in the AMBCA connectome. For example, the AMBCA connectivity matrix exhibited dense intra-regional connectivity within the cortex (blue rectangle) and thalamus (red rectangle). The robust corticothalamic connectivity observed in the AMBCA streamlines (purple rectangle) was preserved in FiberLM-generated matrices but was markedly reduced in those derived from probabilistic tractography.

**Figure 3:**
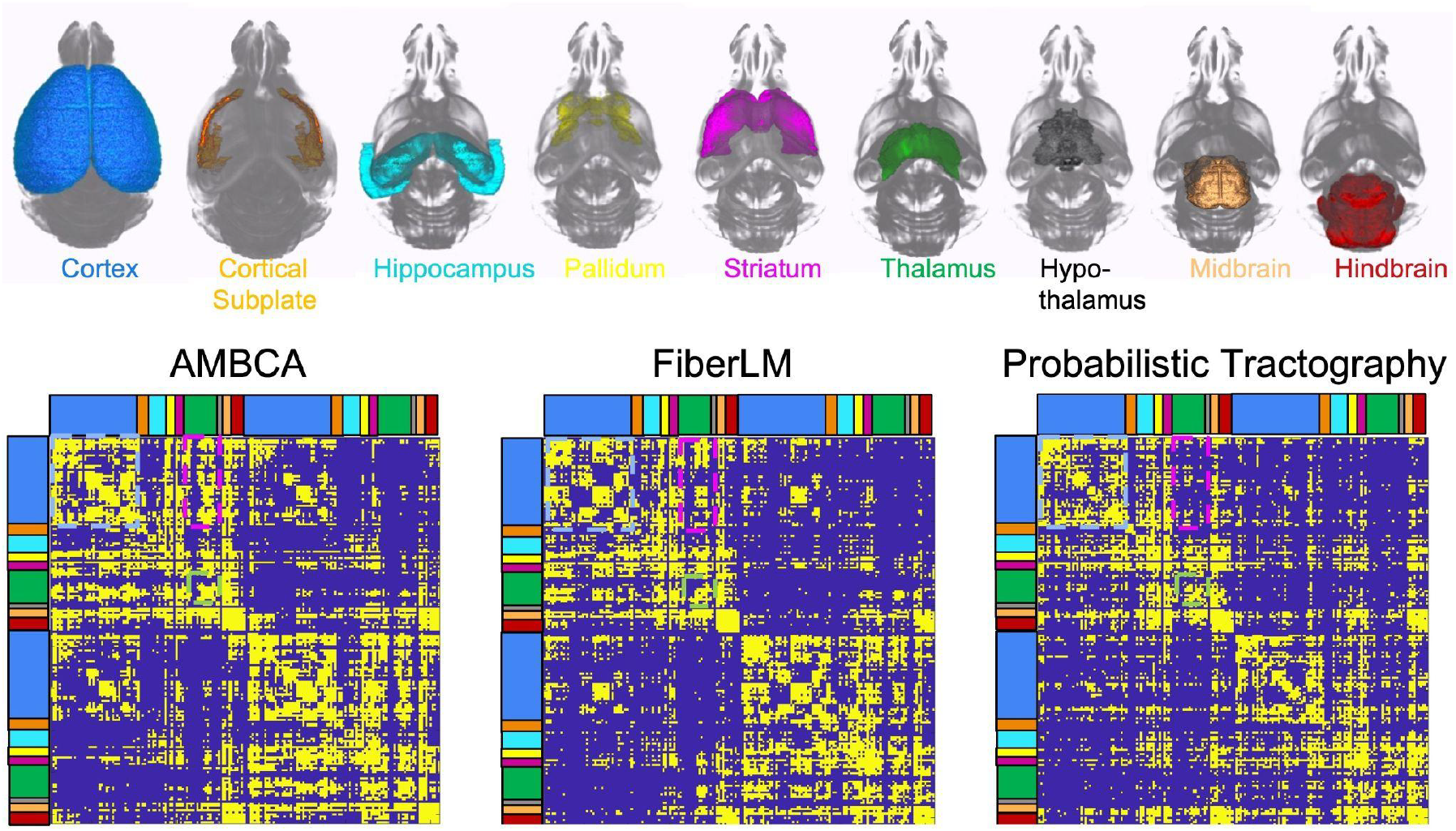
A comparison of whole-brain connectivity matrices based on the tractograms generated using FiberLM and probabilistic tractography. **Top:** Major mouse brain regions used in the analysis were grouped and color-coded. **Bottom:** Binary whole-brain connectivity matrices derived from the Allen reference connectome (left) and a representative mouse brain using FiberLM (middle) and probabilistic tractography (right). The colorbars on the top and left indicate the locations of brain regions in the connectivity matrices.

Graph theory-based network metrics were computed to further characterize whole-brain connectivity across methods (**Fig. 4**). FiberLM-generated brain connectivity matrices exhibited significantly higher global efficiency and clustering coefficient than probabilistic tractography (p = 0.04 and p = 0.047, respectively; n = 6), though both metrics remained lower than those of the AMBCA streamline dataset. Modularity and characteristic path length did not differ significantly between results from FiberLM and conventional tractography. For the four metrics, FiberLM results aligned better with the AMBCA streamlines than probabilistic tractography, suggesting similar network topology.

**Figure 4:**
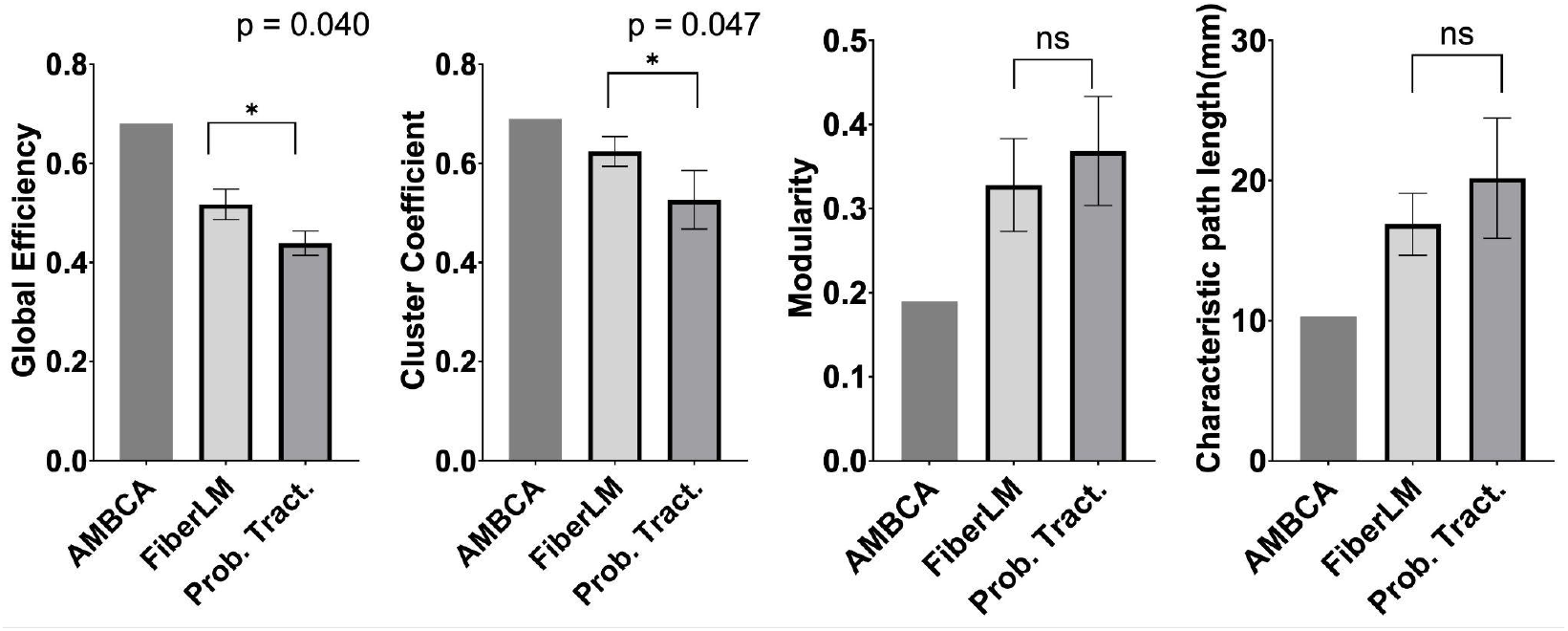
A comparison of network metrics derived from the AMBCA streamline dataset, FiberLM, and probabilistic tractography results (mean ± SEM). * indicates p < 0.05.

In terms of nodal degree (i.e. the number of connections associated with a brain region), FiberLM-generated connectivity matrices exhibited smaller relative differences compared to the AMBCA reference than conventional probabilistic tractography (**Fig. 5A**), with notable differences in the cortical regions. Nodes exhibiting large degree deviations are listed in **Supplementary Table 1**. One example is the connections from the medial group of the dorsal thalamus (MED, **Fig. 5B**). FiberLM-generated streamlines from MED predominantly reached the ipsilateral cortex, with a small number extending to the contralateral cortex and midbrain. In contrast, probabilistic tractography-generated streamlines from MED reached only limited ipsilateral cortical regions with no contralateral cortical projections, but exhibited an excess of streamlines to the midbrain and cerebellum (indicated by the white arrow) compared to the AMBCA reference, resulting in differences in nodal degrees. Another example is the middle corpus callosum (mcc, **Fig. 5C**), a node without significant degree deviation between the two methods. Both FiberLM and probabilistic tractography results showed connectivity comparable to the AMBCA streamlines from the middle corpus callosum. Nevertheless, probabilistic tractography produced an increased incidence of looping and circular streamlines (indicated by the white arrow) not seen in the AMBCA streamlines.

**Figure 5:**
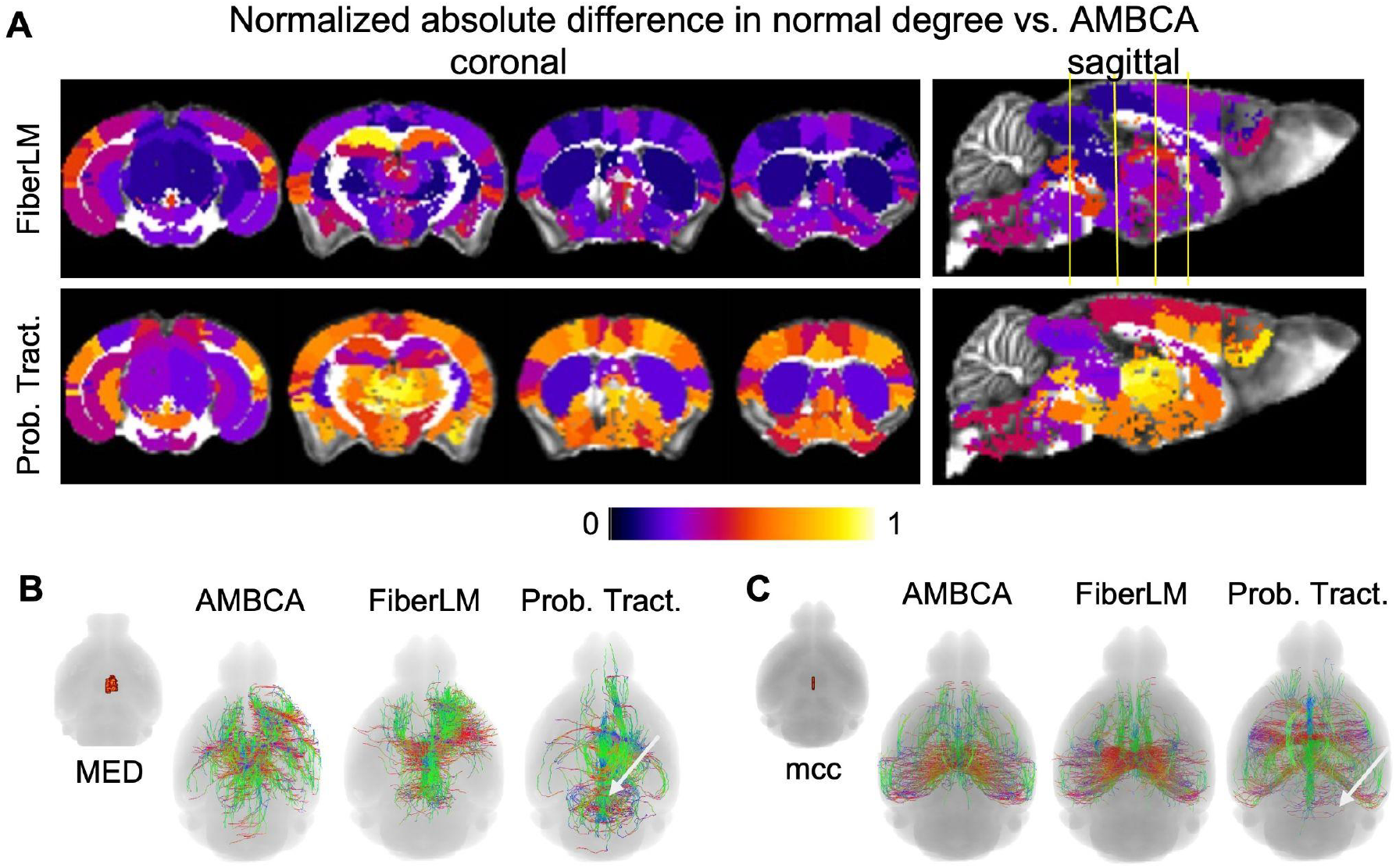
Node-level comparison between FiberLM and probabilistic tractography. (**A**) Regional maps of normalized absolute nodal degree difference for FiberLM and probabilistic tractography with respect to the AMBCA reference. Color intensity reflects the magnitude of relative nodal degree deviation. (**B–C**) Representative streamline reconstructions passing through the medial group of the dorsal thalamus (MED) and the midbody of the corpus callosum, shown for the AMBCA dataset, FiberLM, and probabilistic tractography. (**D**) Mean absolute nodal degree deviation (± SD) across subjects for selected nodes.

### 3.3 FiberLM reduced false positive and false negative connections

Using the AMBCA streamlines as the ground truth, probabilistic tractography produced a substantial number of false positive connections (**Fig. 6A**). Clusters of spurious connections were distributed between cortical subregions and between cortical and thalamic regions as well as across hemispheres. By contrast, false positive connections in FiberLM results were considerably reduced and became more isolated. **Fig. 6B** shows two representative false positive connections. Between the lateral septal complex (LSX) and secondary somatosensory area (SSs), probabilistic tractography generated diffuse streamlines spanning broad cortical and subcortical territories, whereas FiberLM produced only a small number of spatially confined streamlines. Similarly, between the frontal pole (FRP) and motor-related pons (Pons), probabilistic tractography produced a large streamline bundle, while FiberLM returned only a few streamlines. The small number of false positive streamlines generated by FiberLM can be easily removed in the connectivity matrix by thresholding. It should be noted that the Allen tracer reference is neither exhaustive nor uniform in its anatomical coverage, as it is constrained by the availability of tracer injections; consequently, some false positives may reflect connections that are currently unverified rather than genuinely absent. In the whole brain maps of regional false-positive rates (**Fig. 6C**), probabilistic tractography yielded notably higher rates in cortical, hippocampal, and thalamic regions than FiberLM. Overall, FiberLM produced a significantly lower false-positive rate (7.7 ± 2%) than conventional probabilistic tractography (14.3 ± 6%; p < 0.01, paired t-test).

**Figure 6.**
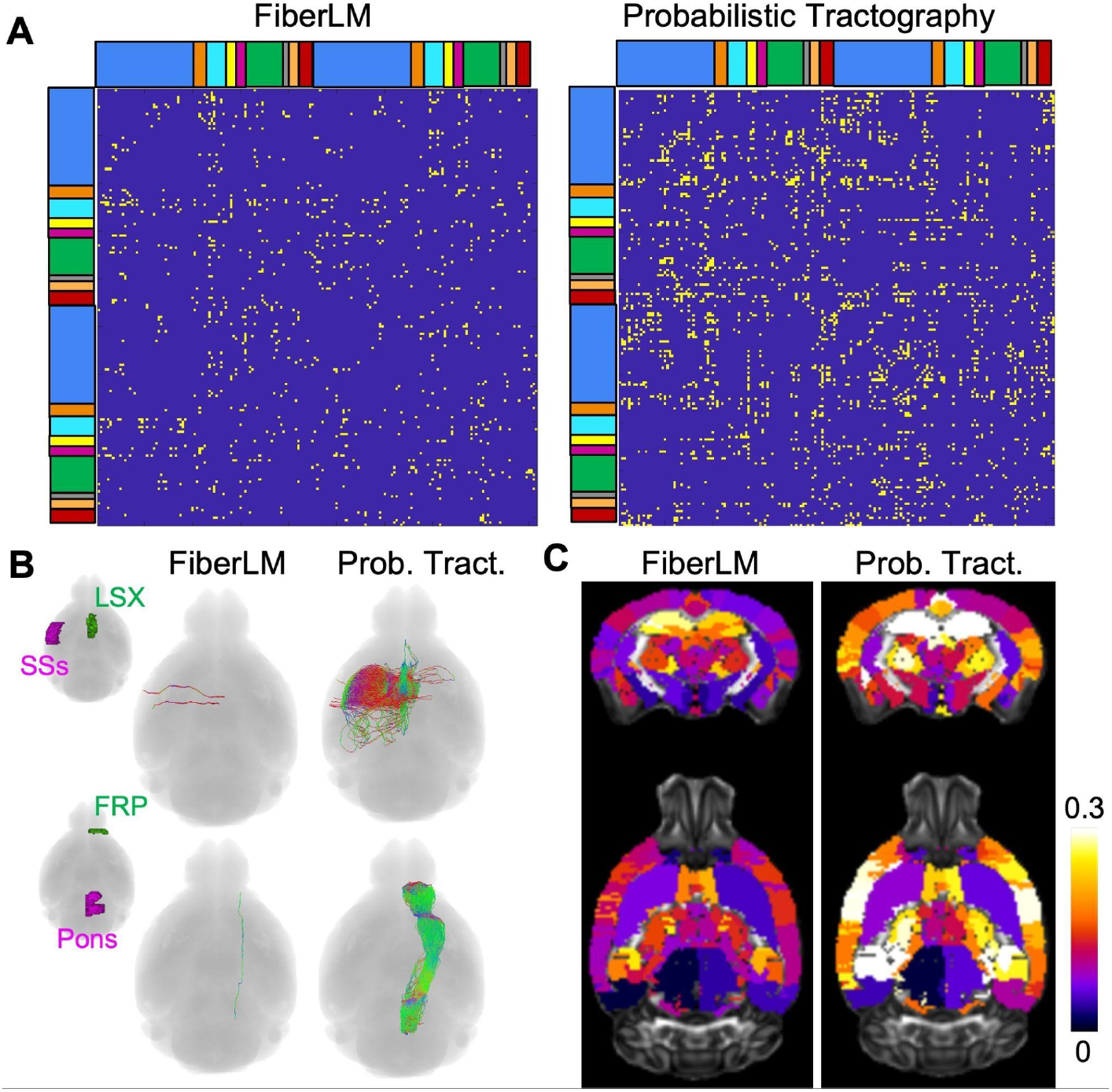
False-positive connections generated by FiberLM and probabilistic tractography. (**A**) An overview of false-positive connections in the connectivity matrices from FiberLM (left) and probabilistic tractography (right). (**B**) Representative examples of false-positive connections generated by probabilistic tractography between the lateral septal complex (LSX) and the secondary somatosensory area (SSs) and between the frontal pole (FRP) and the motor related pons (Pons). (**C**) Maps of regional false-positive rates for FiberLM and probabilistic tractography.

The FiberLM results also contained less false negative connections than the results from conventional probabilistic tractography (**Fig. 7A**), particularly along cortico-thalamic and cortico-brainstem connections. However, a number of false negatives still persisted for FiberLM for connections involving the hippocampus and adjacent retrosplenial cortex. **Fig. 7B** illustrates the spatial distribution of false-negative streamlines. In the whole brain maps of regional false-negative rates (**Fig. 7C**), probabilistic tractography yielded notably higher rates in cortical and thalamic regions than FiberLM, but lower rates in the hippocampus. Overall, FiberLM achieved a significantly lower false-negative rate (23 ± 2%) compared with probabilistic tractography (30 ± 1%; p < 0.01, paired t-test).

**Figure 7.**
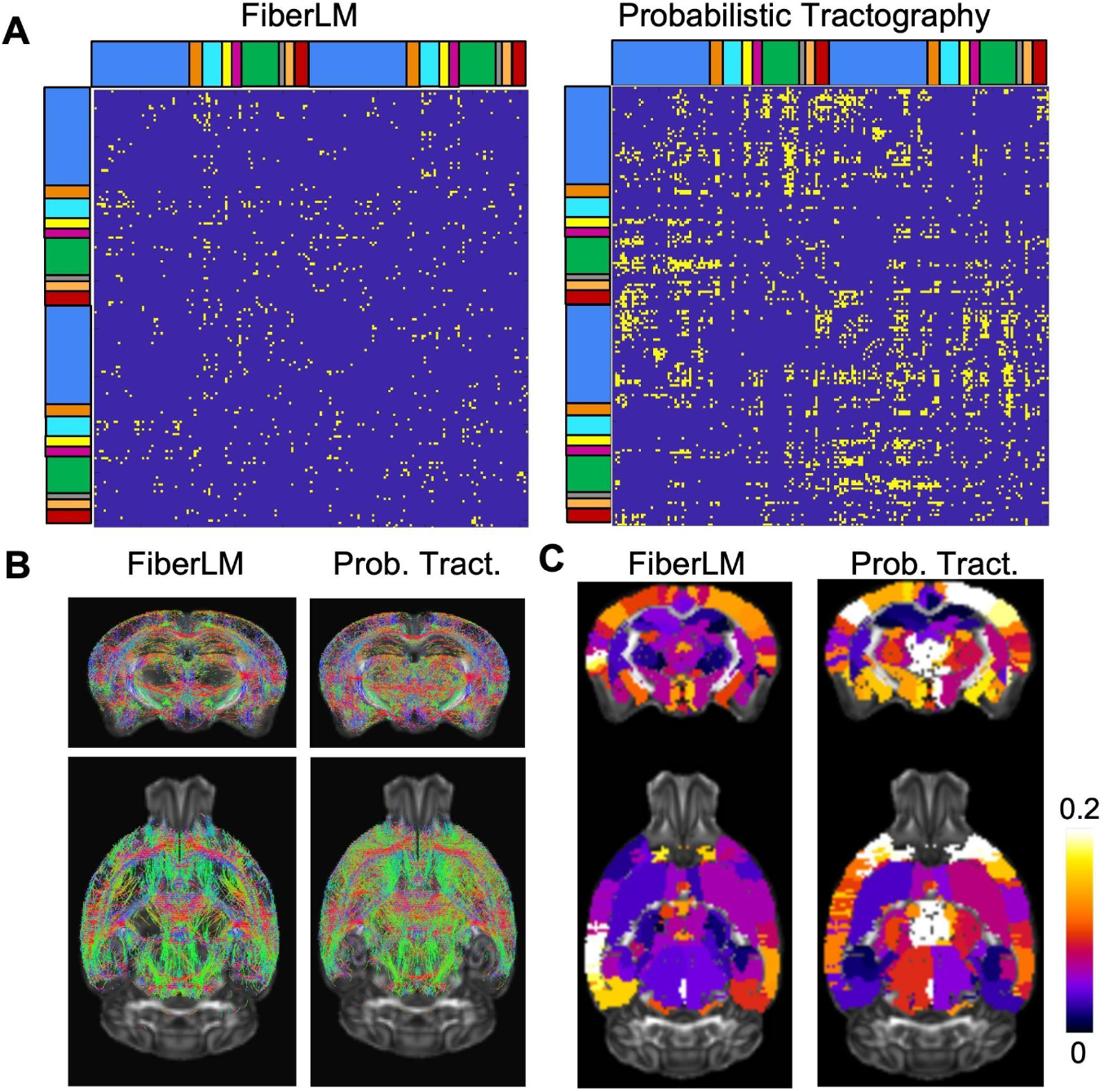
False-negative connections by FiberLM and probabilistic tractography. (**A**) An overview of false negative connections in the whole brain connectivity matrices from FiberLM and probabilistic tractography. (**B**) false-negative connections displayed using their corresponding streamlines from the AMBCA dataset. (**C**) Maps of regional false-negative rates for FiberLM and probabilistic tractography.

Among true positive connections, FiberLM results showed better spatial agreements with the AMBCA streamlines than the probabilistic tractography results. For example, both probabilistic tractography and FiberLM found connections between the retrosplenial area ventral (RSPv) and reticular nucleus (RT) (**Fig. 8A**) and between presubiculum (PRE) and pons (ROI) (**Fig. 8B**). The streamlines generated by probabilistic tractography contained more spurious streamlines than the FiberLM results. Similar cases were also observed for the connections between the orbital area lateral (ORBl) and primary somatomotor area (Mop) (**Fig. 8C**) and between the somatosensory area upper limb (SSp-ul) and caudoputamen (CP) (**Fig. 8D**). Overall, FiberLM results showed better spatial agreement with the AMBCA streamlines than probabilistic tractography (Dice score: 0.57±0.1 vs. 0.21 ±0.12, p<0.001, for conventional probabilistic tractography).

**Figure 8.**
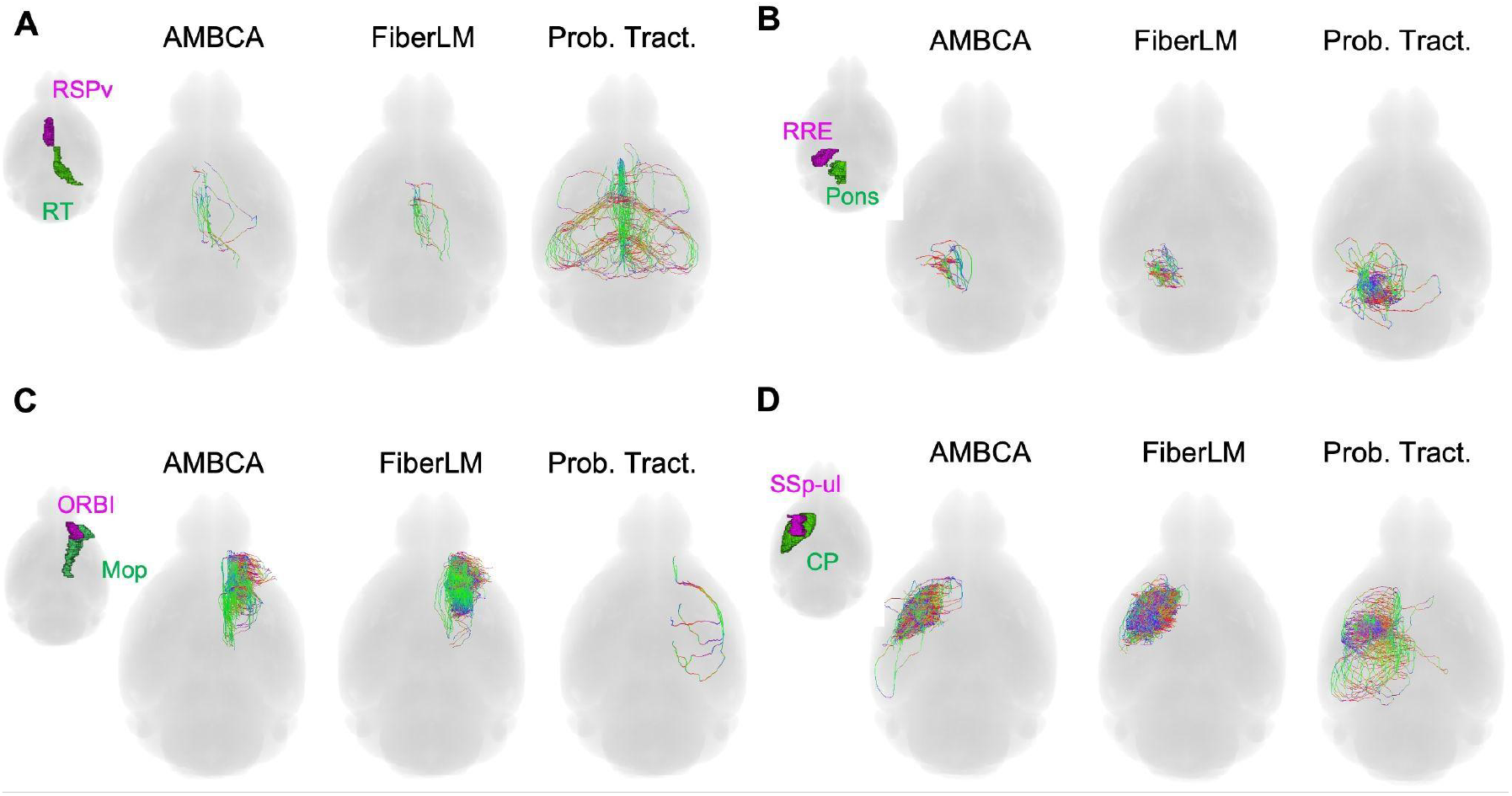
Comparisons between probabilistic tractography and FiberLM for selected true positive connections.

### 3.4 Attention map

Voxel-wise attention maps (**Fig. 9A**) demonstrate that attention was heterogeneously distributed across the brain, with elevated values observed in subcortical and peri-thalamic regions, while large, smoothly organized white-matter bundles show comparatively moderate attention levels. The spatial pattern of attention resembled the distribution of local AMBCA streamline curvature, with regions of high geometric complexity exhibiting stronger attention allocation. Further analysis (**Fig. 9B**) showed a significant positive association between voxel-wise local curvature and attention (Spearman R = 0.46, p < 0.0001). The density plot reveals a monotonic increase in the binned mean attention as log-curvature increases, indicating that attention systematically scales with geometric complexity. Notably, this relationship is not purely linear: low-curvature regions exhibited consistently low attention values, whereas a subset of high-curvature voxels displayed a pronounced upward trend in mean attention. Taken together, these findings suggest that the network preferentially weighted voxels characterized by greater local bending, consistent with selective attention engagement at geometrically complex regions where streamline propagation is inherently ambiguous.

**Fig. 9:**
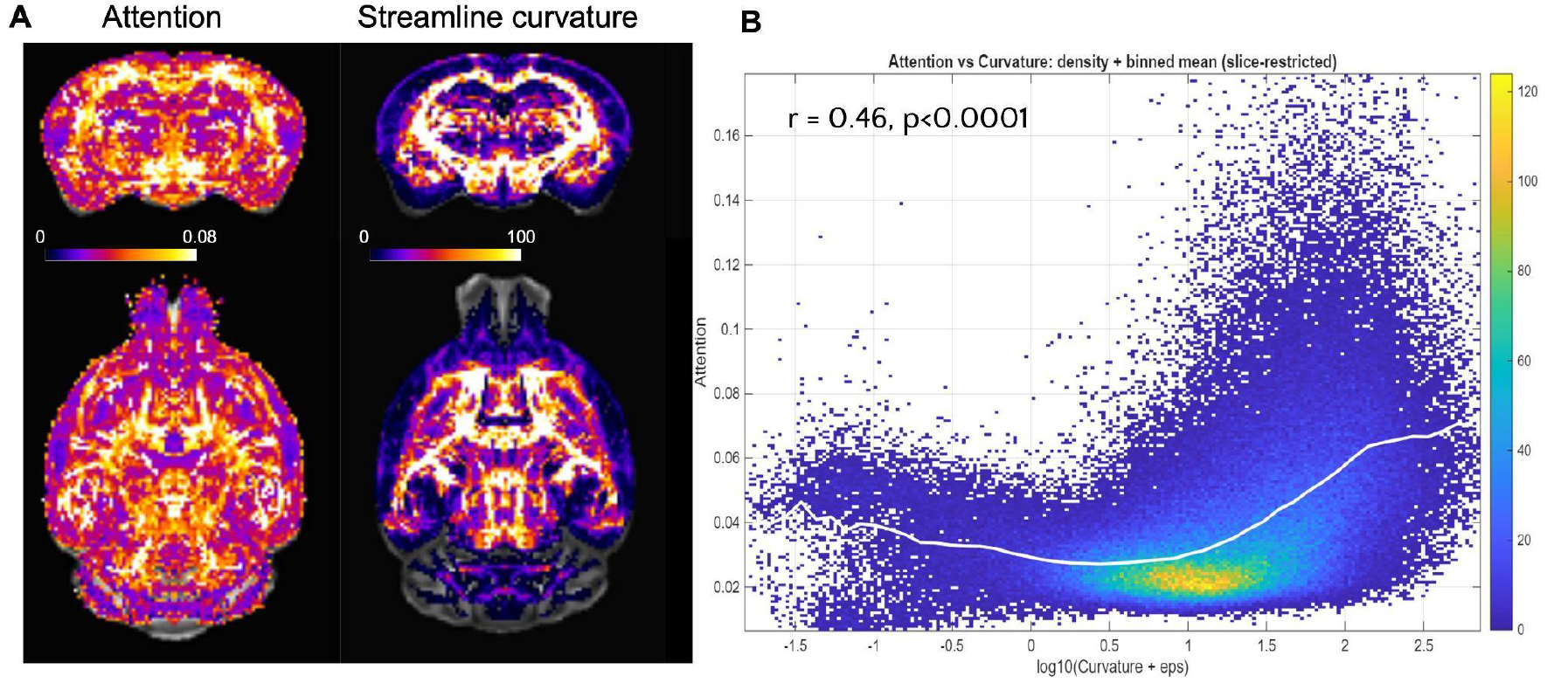
Association between local streamline curvature and Transformer attention. (**A**) Axial and horizontal maps of averaged attention and local mean curvature of AMBCA streamline segments in the mouse brain. (**B**) Voxel-wise density plot of ln(curvature) versus attention (slice-restricted). The white curve indicates the mean attention at different curvatures. A significant positive association is observed (Spearman R = 0.46, p < 0.0001), indicating that attention generally increases with local streamline curvatures.

## Discussion

Our findings demonstrate that a Transformer-based model trained using the AMBCA streamlines can substantially improve tractography accuracy. Prior work has explored using histological ground truth to improve tractography, including effort by Schilling et al. (Schilling et al., 2018) in nonhuman primate brains and Liang et al. (Liang et al., 2023) in the mouse brain, though these studies focused primarily on improving fODF estimation. The present work extended the deep learning approach to the streamline generation step, leveraging a transformer-based architecture to learn directly from tracer-derived axonal trajectories. The reductions in both false positives and false negatives suggest that the model can reconstruct the majority of macroscopic connections in the mouse brain with anatomically accurate spatial trajectories, a prerequisite for reliable connectivity-based analyses.

### 4.1 FiberLM utilizes anatomical information encoded in the AMBCA streamline data

Previous studies have used dMRI tractography to reconstruct the whole mouse brain connectome, but validation of the reconstructed network remains difficult, with most efforts focusing on a limited number of tracts. The mouse brain poses particular challenges, as the majority of its brain volume is gray matter, whose microstructural organization is more complex than white matter. In gray matter, the estimated fFODs often show multiple potential directions for streamline propagation, and conventional probabilistic tractography struggles in the gray matter because it mostly relies on local information to determine whether a streamline should follow a particular direction, leading to potential false-positive and false-negative connections. For example, the connections between the cortex and thalamus are critical for brain function given the thalamus’s role as the brain’s primary relay station and the cortex’s role in higher-order processing. In the mouse brain, these connections must course through densely packed fibers in the external capsule or traverse the striatum. Conventional probabilistic tractography has been shown to generate multiple false-negative connections in these regions (Arefin et al., 2023; Aydogan et al., 2018; Grisot et al., 2021).

One strategy to address these challenges is to incorporate anatomical constraints directly into the tractography process, either through predefined inclusion or exclusion regions (Smith et al., 2012) or through clustering/classification in physical or latent spaces (Dumais et al., 2023). While effective for well-characterized pathways, this approach is inherently limited by the extent of existing anatomical knowledge and is less suitable for complex or poorly understood connections. In this study, we leveraged the AMBCA streamline dataset to improve streamline propagation in a data-driven manner, covering a broader range of connections and their trajectories than what could be captured by manually defined constraints. Rather than imposing hard rules derived from incomplete prior knowledge, we hypothesize that FiberLM learned the natural coursing patterns of streamlines through the brain from this rich dataset, enabling it to make fewer erroneous turns and direct streamlines toward anatomically correct destinations rather than spurious ones. This is supported by FiberLM’s ability to reconstruct many corticothalamic connections that conventional probabilistic tractography failed to recover (e.g. **Fig. 5B**), suggesting that the model internalized the complex routing rules governing these pathways.

### 4.2 Transformer-based contextual modeling enhances global coherence

The adoption of the Transformer architecture for tractography was inspired by its success in large language models, where sequential tokens are interpreted within a broader context. In tractography, a 3D streamline trajectory is analogous to a sentence, with the underlying anatomy serving as the context that governs how each step should be interpreted. A key advantage of this architecture is its ability to capture long-range dependencies along streamline trajectories. Unlike RNN-based models, which propagate local memory sequentially and are therefore limited in their ability to integrate distant contextual information, the self-attention mechanism evaluates the entire trajectory history simultaneously and dynamically weights the relevance of each step. This property is reflected in whole-brain comparisons (**Figs. 5 and 8**), where FiberLM produced smoother, anatomically continuous tracts across crossing-fiber regions such as the internal capsule and midbrain — areas where conventional probabilistic tractography tended to fragment or deviate. Consistent with this, Yang et al. (Yang et al., 2025) reported that the attention-based formulation reduced curvature errors and helped maintain correct propagation paths through fiber-crossing regions in the human brain. In the mouse brain, FiberLM similarly exhibited lower false-positive density in the cortico-subcortical projections (cortex→thalamus and cortex→midbrain; **Fig. 7D**), while preserving valid interhemispheric and hippocampal pathways. Together, these findings suggest that attention weighting improves anatomical coherence by integrating local orientation cues with global trajectory history, without sacrificing sensitivity.

A natural question is whether FiberLM simply memorized the trajectories present in the AMBCA training data rather than learning generalizable tracking rules. This is unlikely for several reasons. First, the inputs to the model were FODs without anatomical locations of the corresponding streamlines. Second, the model was evaluated on dMRI data from a separate cohort of mice not seen during training, and performance remained consistent. Third, tractography results in the cerebellum, a region with only sparse streamline coverage in the AMBCA dataset, still showing dense streamlines, indicating that the model is capable of generating plausible trajectories beyond what was directly observed during training. These observations suggest that FiberLM learned underlying anatomical principles of axonal pathways rather than reproducing memorized streamline patterns.

Beyond reconstruction accuracy, the learned attention maps reveal structured, biologically meaningful patterns (**Fig. 9A**) and provide a window into which anatomical regions most influence model decisions during streamline propagation. Elevated attention is consistently observed in subcortical projection systems, particularly within thalamic and ventral diencephalic regions. This spatial selectivity suggests that the Transformer preferentially emphasizes regions characterized by curved pathways and pathway convergence rather than tract volume or streamline density. Such regions likely represent decision-critical bottlenecks where propagation requires integration of broader contextual information, supporting the interpretation that attention reflects complexity of axonal pathways in a region. This observation hints that, for future model refinement, prioritizing axonal pathways that pass through the high attention regions may enhance tractography.

### 4.4. Limitations and considerations

Several caveats should be noted. First, this work assumes that structural connectivity in the inbred adult C57BL/6 mouse brain is highly congruent at the macroscopic and mesoscopic levels, such that AMBCA streamlines can supervise model training using dMRI data from a separate cohort of age-matched mice. While tractography results of major white matter tracts (**Fig. 2**) support this assumption, it may not hold for smaller tracts, and additional validation is needed. Second, although the AMBCA provides unparalleled mesoscale tracer coverage (>2,000 injections), it lacks individual-level whole-brain ground truth. Consequently, FiberLM learned from aggregated population priors, which may obscure individual variability and hemispheric asymmetry. Regional imbalance in tracer injection density, particularly in the cerebellum, brainstem, and olfactory areas, may further limit model generalization in these regions. Additionally, the non-uniform spatial sampling of tracer streamlines means that streamline density does not equate to true axonal counts and therefore cannot be interpreted as absolute connectivity strength. While our whole-brain blind validation minimizes overfitting, additional experiments incorporating tracer subsets or cross-cohort testing will be needed to fully establish model generalizability.

Additional limitations relate to the scope of the training data. The current implementation is focused on the normal mouse brain, and extending FiberLM to disease models will require further validation. The model relies on FODs estimated from dMRI data using the CSD algorithm. As previously reported, CSD-derived FODs can deviate from the true orientation of axonal projections in several hippocampal and cortical regions due to the presence of dense dendritic structures, which may explain the higher false-negative rates observed in the hippocampus (**Fig. 7C**) (Wu and Zhang, 2016; Liang et al., 2023). Incorporating more accurate FOD estimates, such as those derived from deep learning models trained on histological data (Schilling et al., 2018; Nath et al., 2019; Liang et al., 2023; Zhu et al., 2025), will likely further improve tractography outcomes. Finally, the current implementation requires dMRI data to be spatially normalized to the ARA space in order to leverage anatomical information, adding an additional preprocessing step that may pose challenges for brains with severe deformation or pathological changes such as tumors.

## Conclusion

In summary, FiberLM improved the spatial agreement between mouse brain dMRI tractography results and AMBCA streamlines. In particular, FiberLM reduced false-positive and false-negative connections compared to conventional probabilistic tractography. Furthermore, the model’s attention maps offered interpretable insights into the critical junctions along axonal trajectories that most significantly influence the tractography procedure. Our results demonstrate the feasibility of using viral-tracer-based connectivity data to guide dMRI tractography.

## Data Availability

The data that support the findings of this study are available on request from the corresponding author. The data is not publicly available due to privacy or ethical restrictions.

## Author Contributions

Conceptualization: Z.L., J.Z. Data acquisition: Z.L., J.Z. Formal analysis: R.W., Z.L., J.Z. Writing—original draft: R.W., Z.L., J.Z. Writing—review and editing: R.W., Z.L., J.Z.

## Ethics

All animal procedures were conducted in accordance with protocols approved by the Institutional Animal Care and Use Committee (IACUC) at New York University Grossman School of Medicine.

## Acknowledgement

This study was supported by the National Institute of Health R01NS102904, R01HD074593, U24NS135568, and RM1MH138309. MR imaging was performed at the Preclinical Imaging Center supported by 1S10OD018337-01 and Center of Advanced Imaging Innovation and Research (CAI2R, www.cai2r.net), a Biomedical Technology Resource Center supported by NIBIB with the award P41 EB017183.

## Supplimentary

**Table S1:**
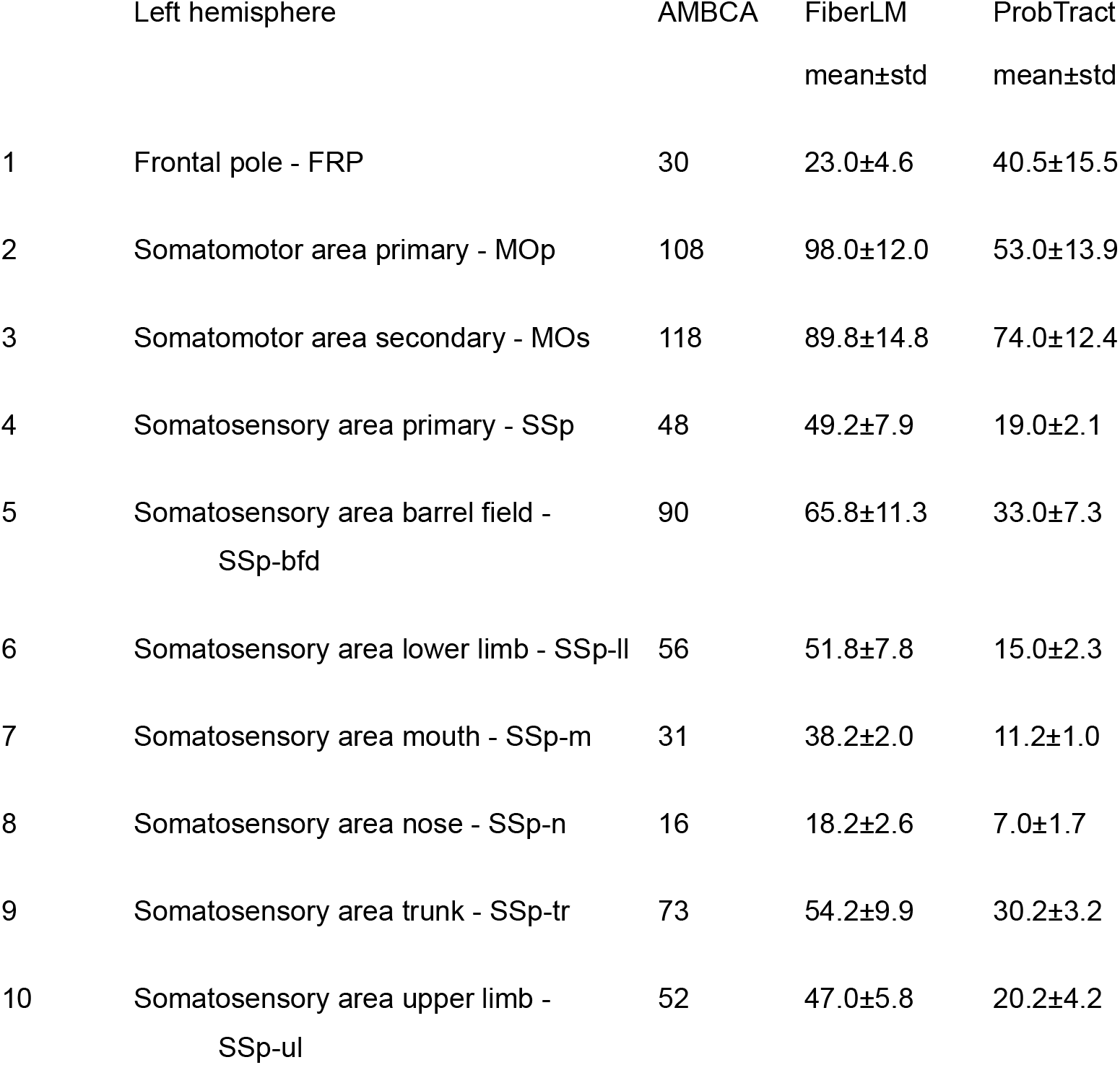
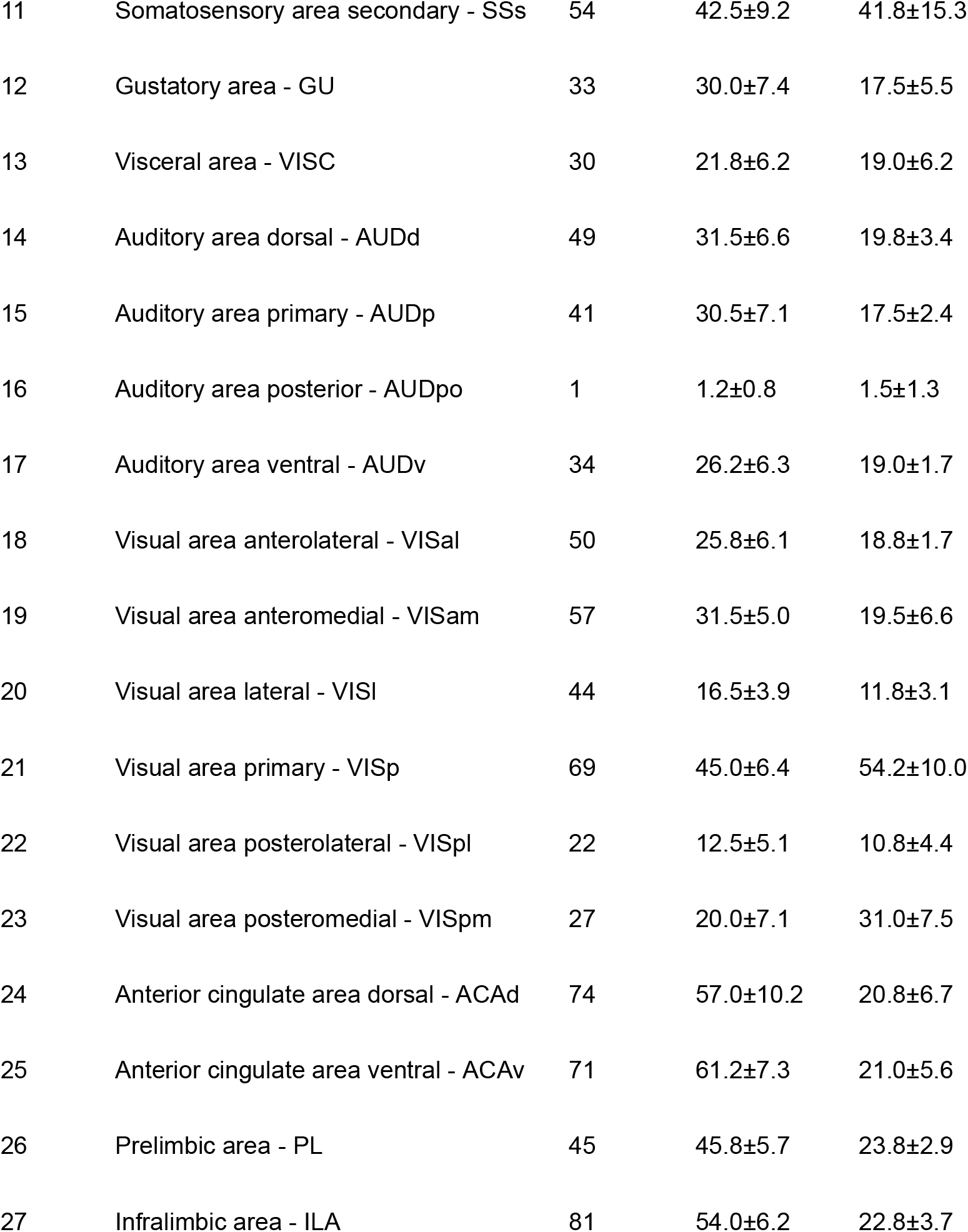
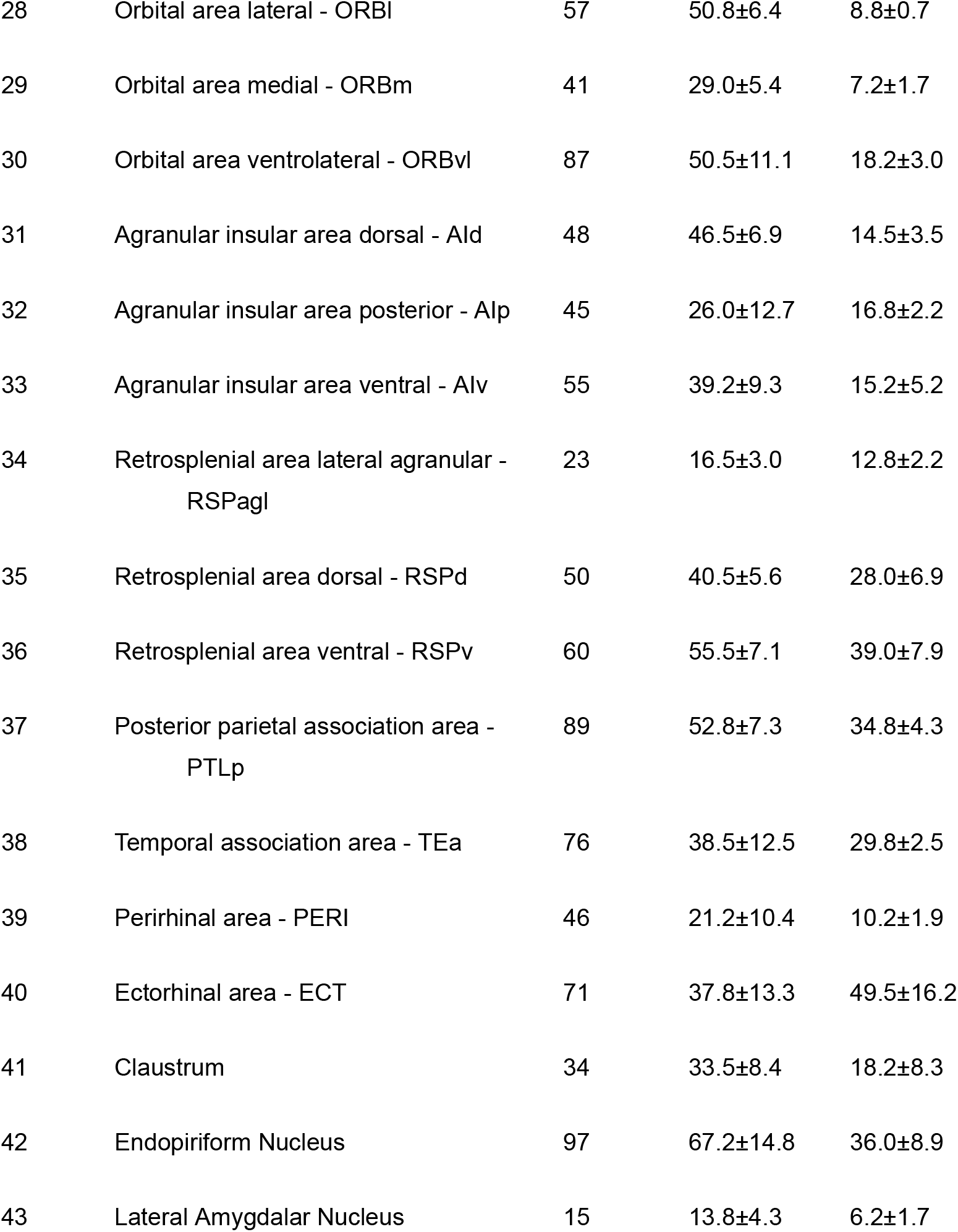
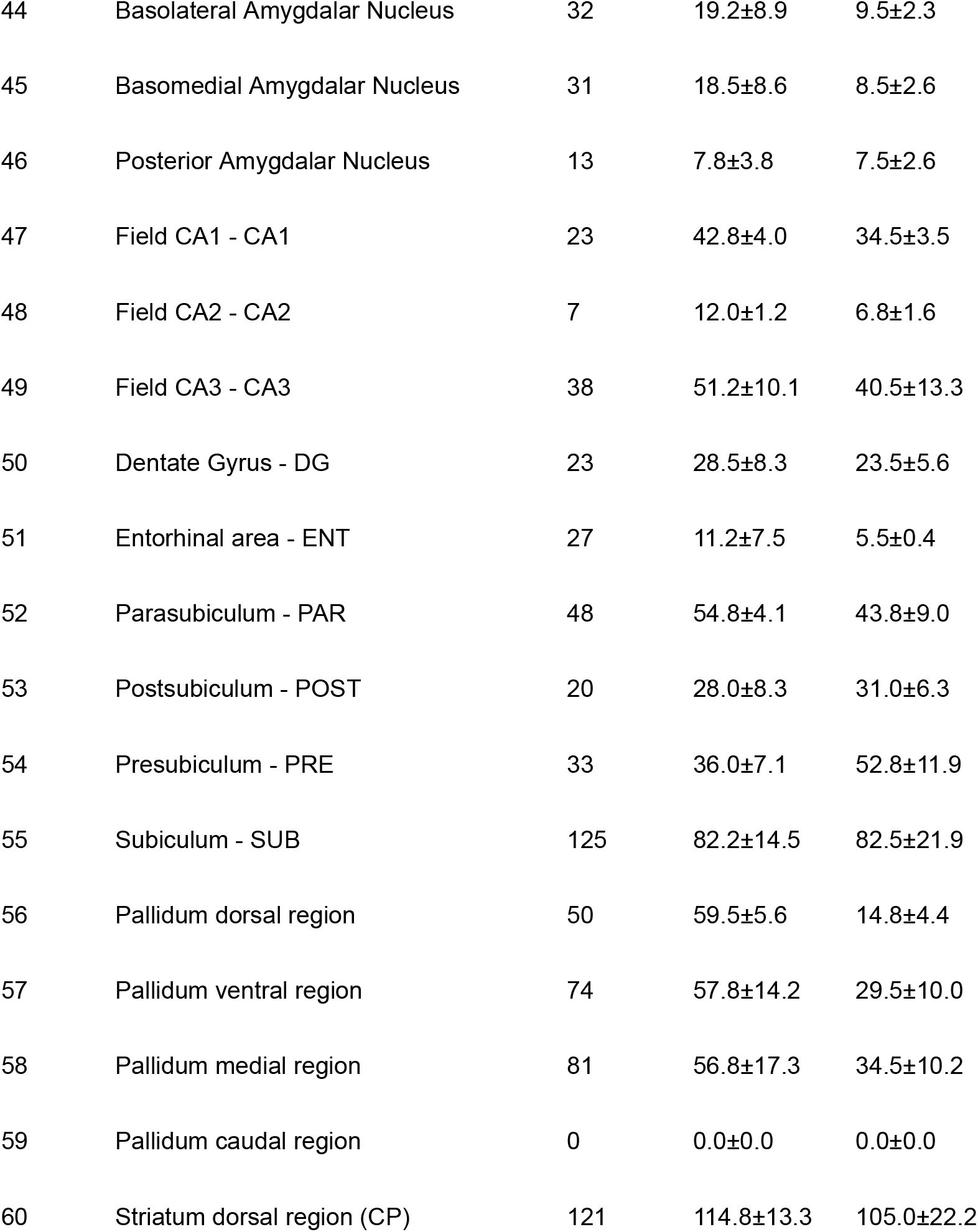
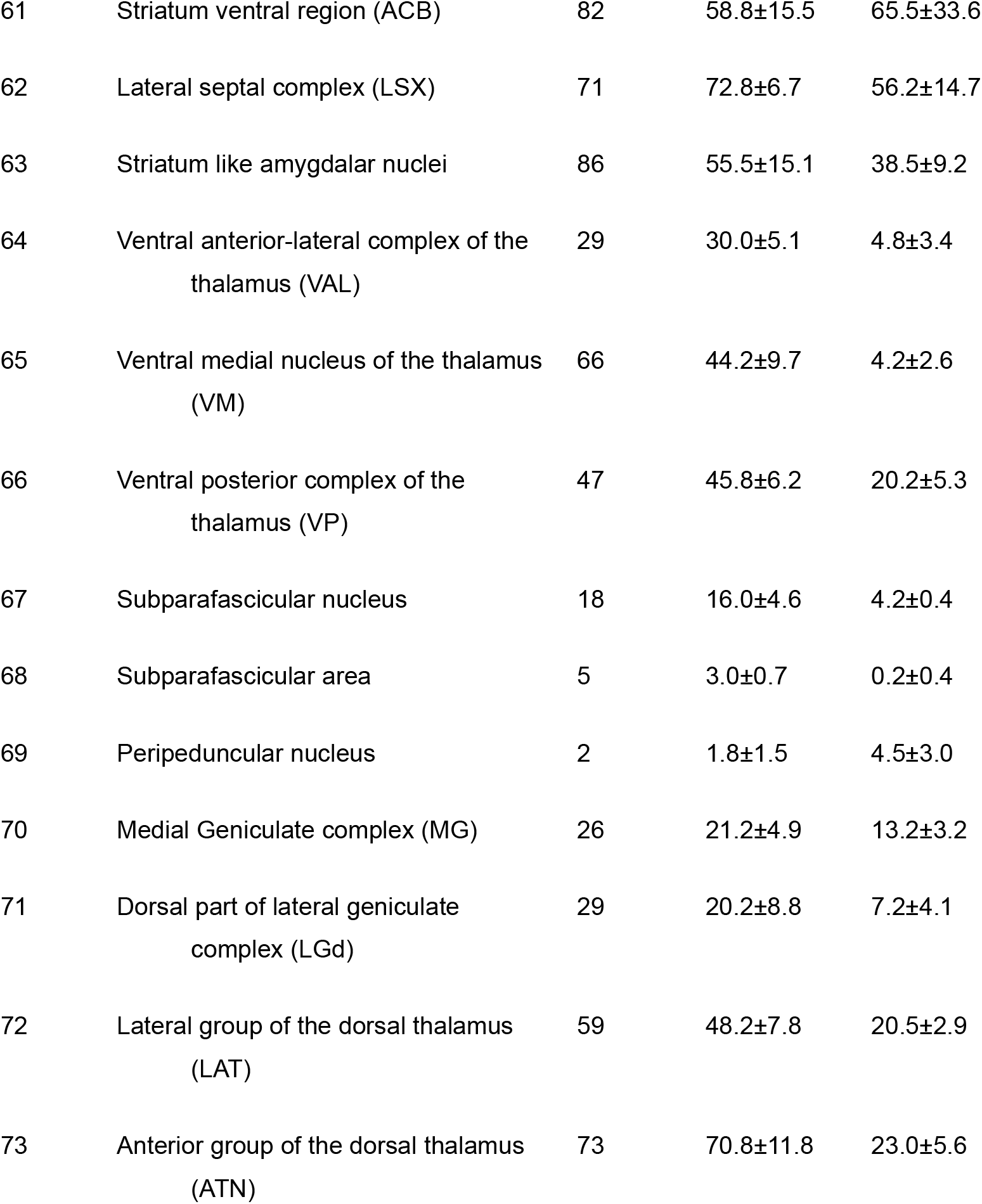
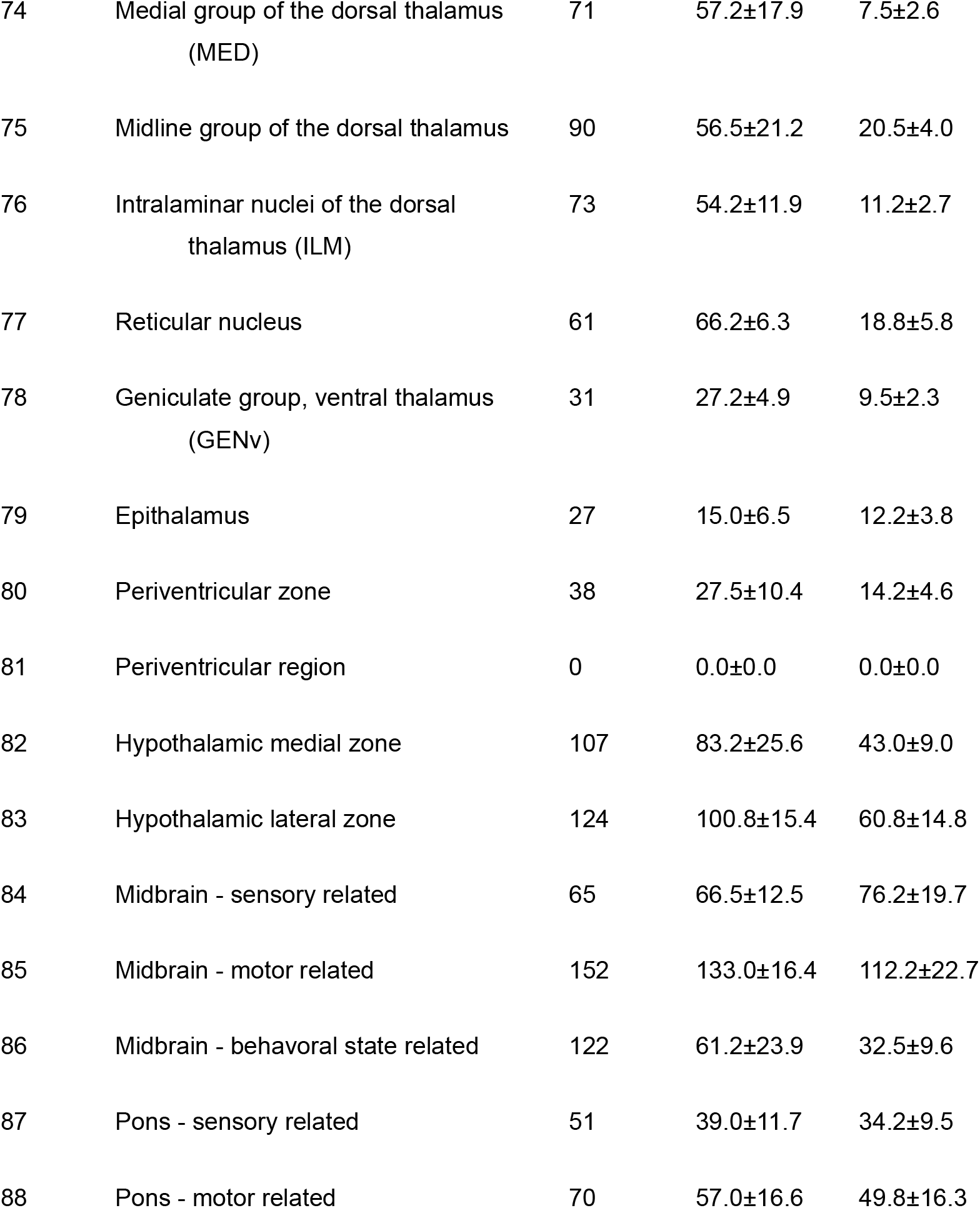
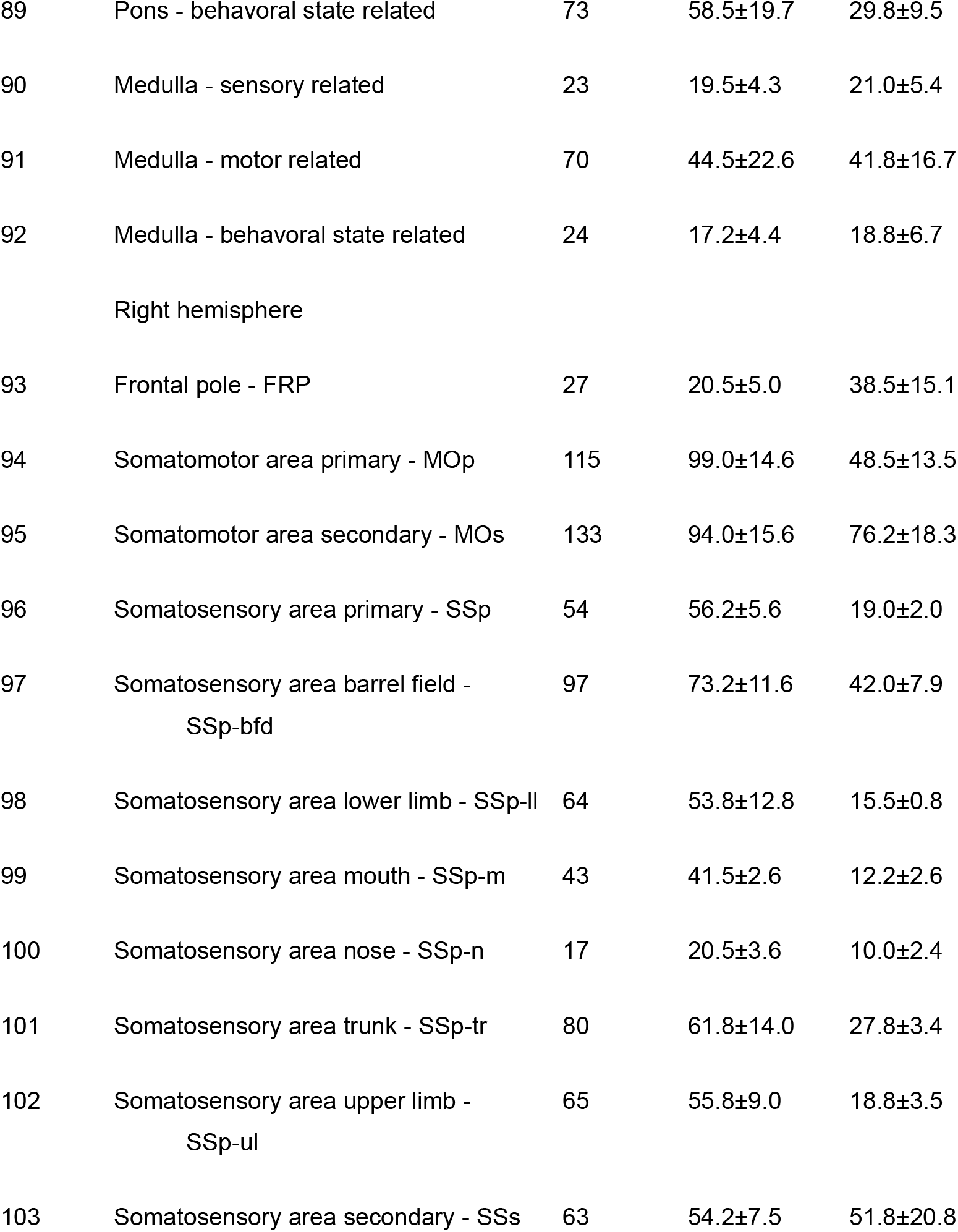
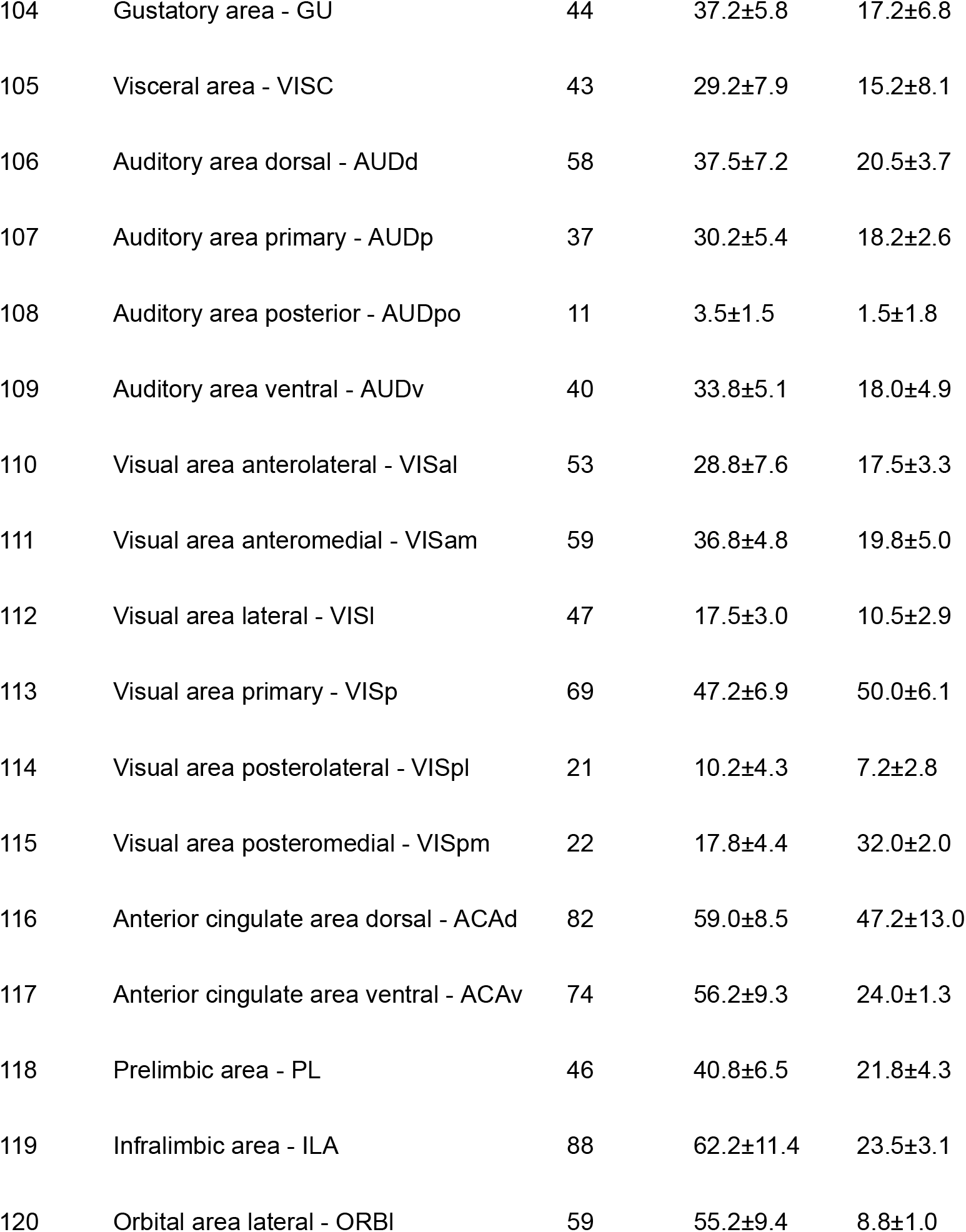
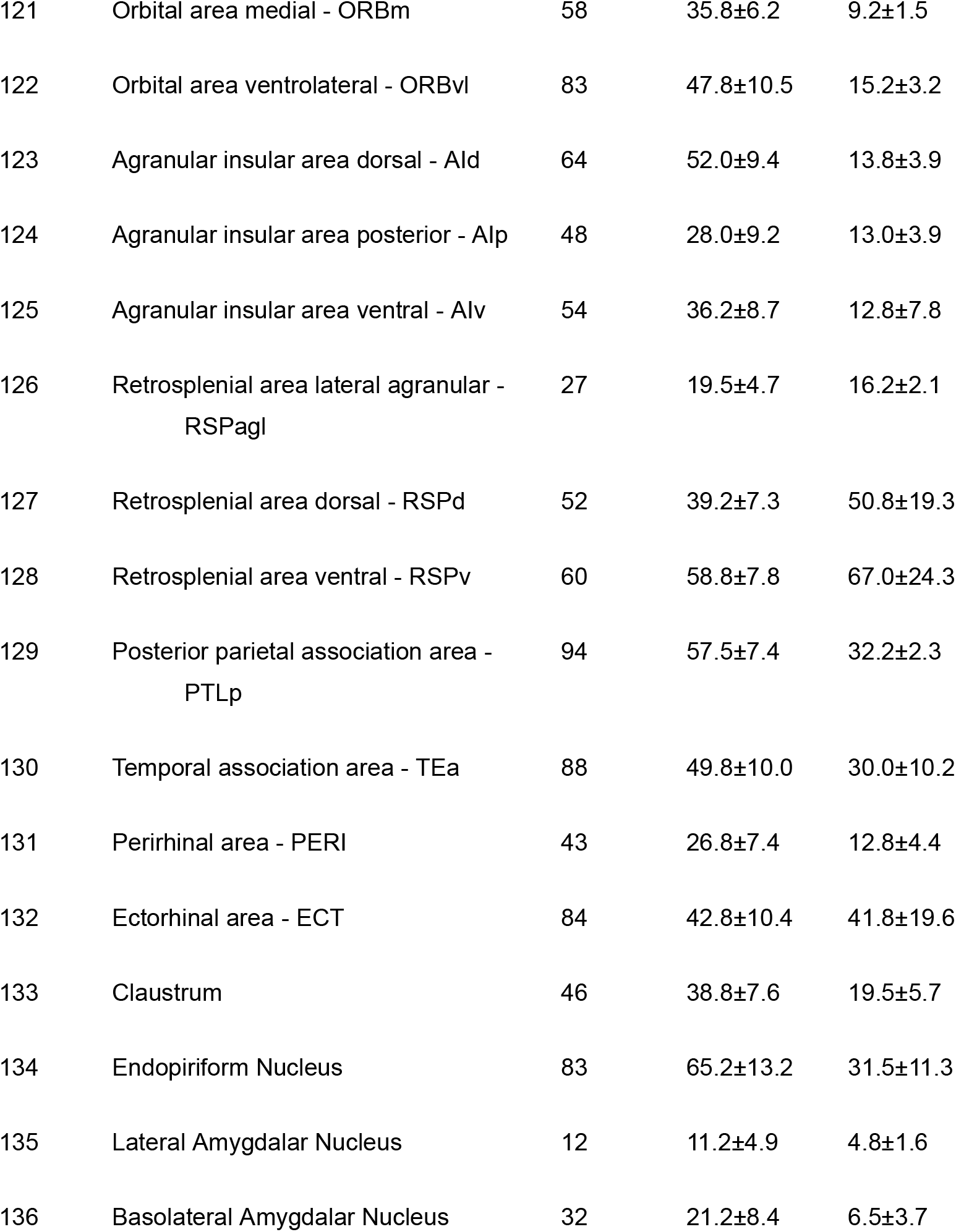
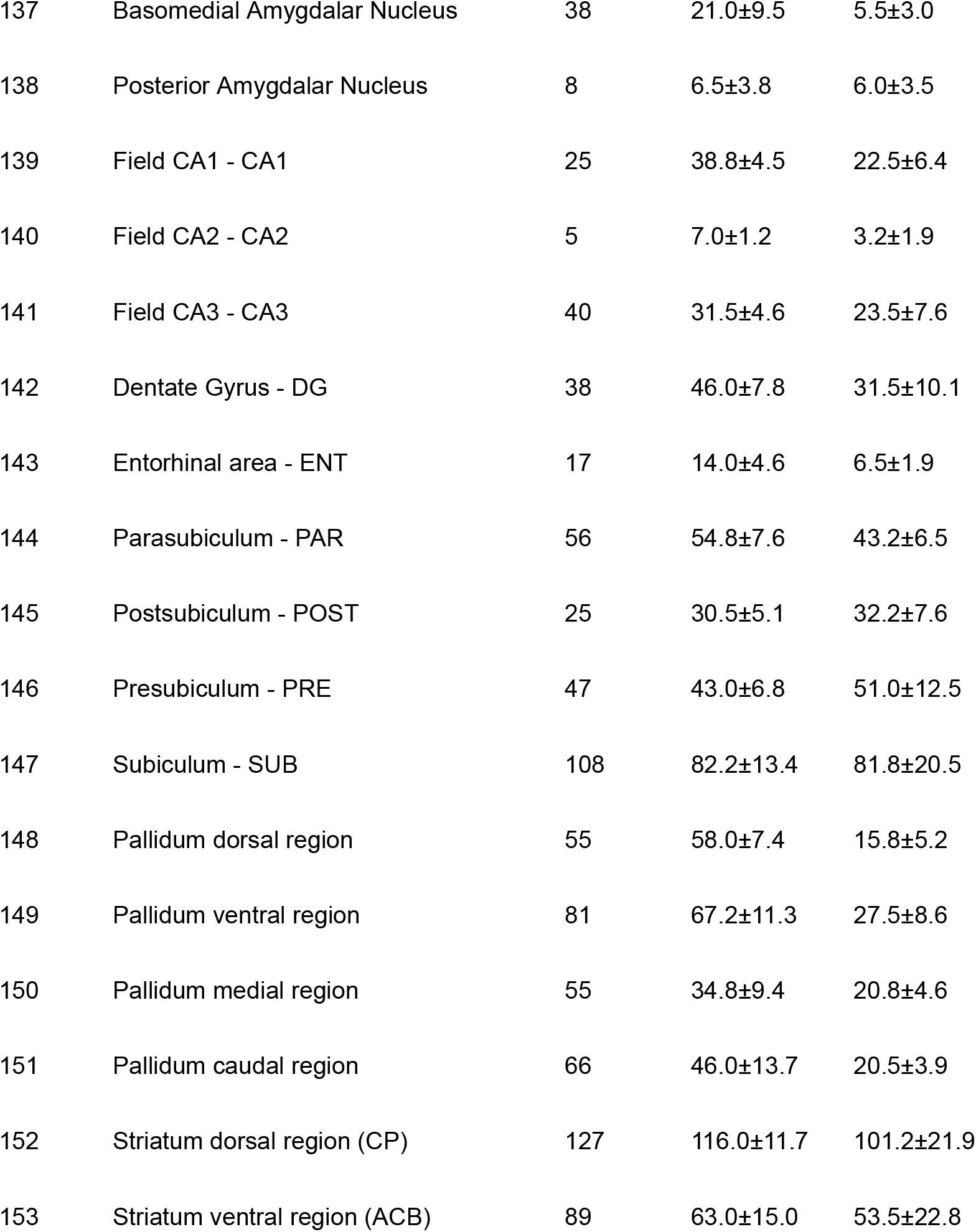
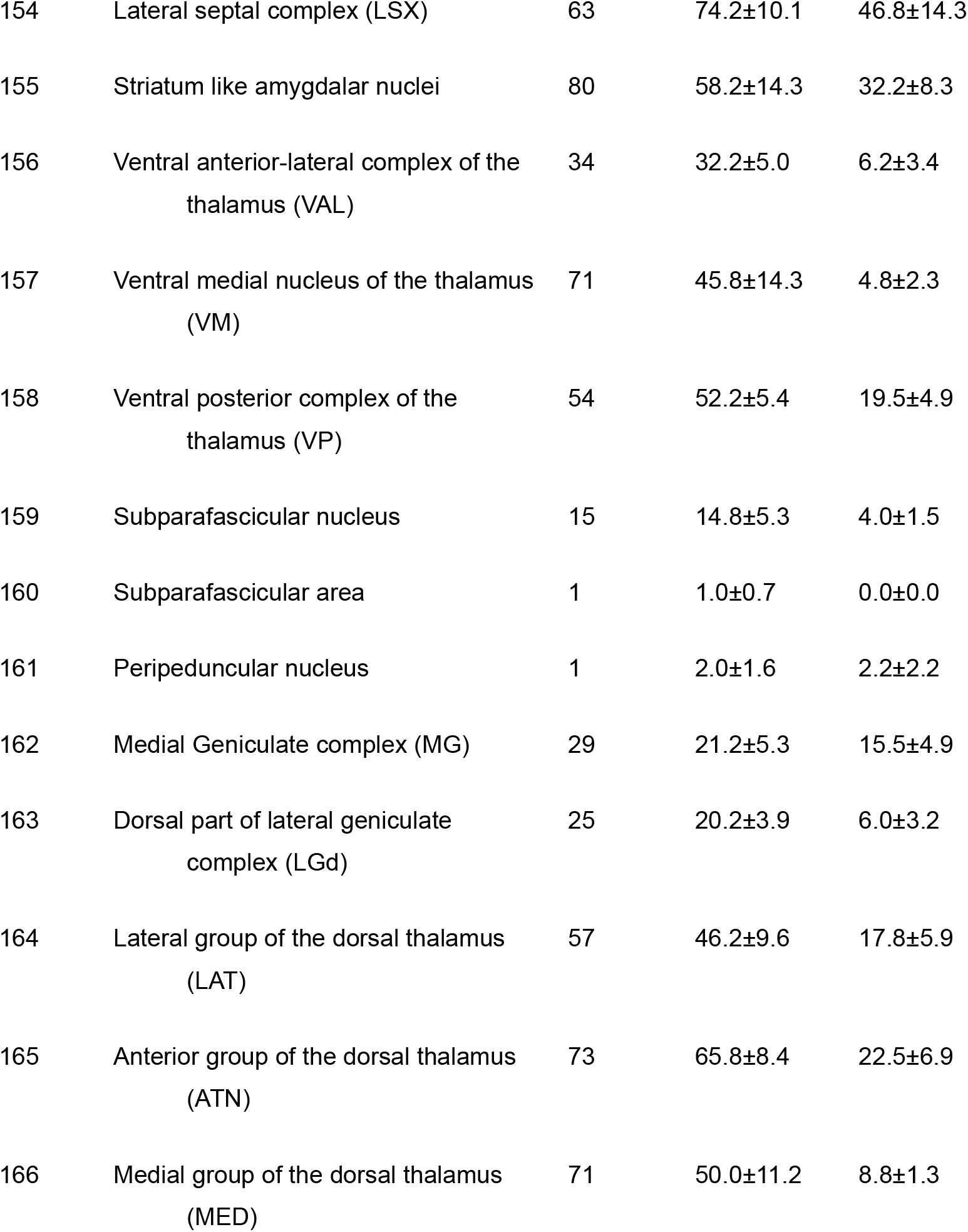
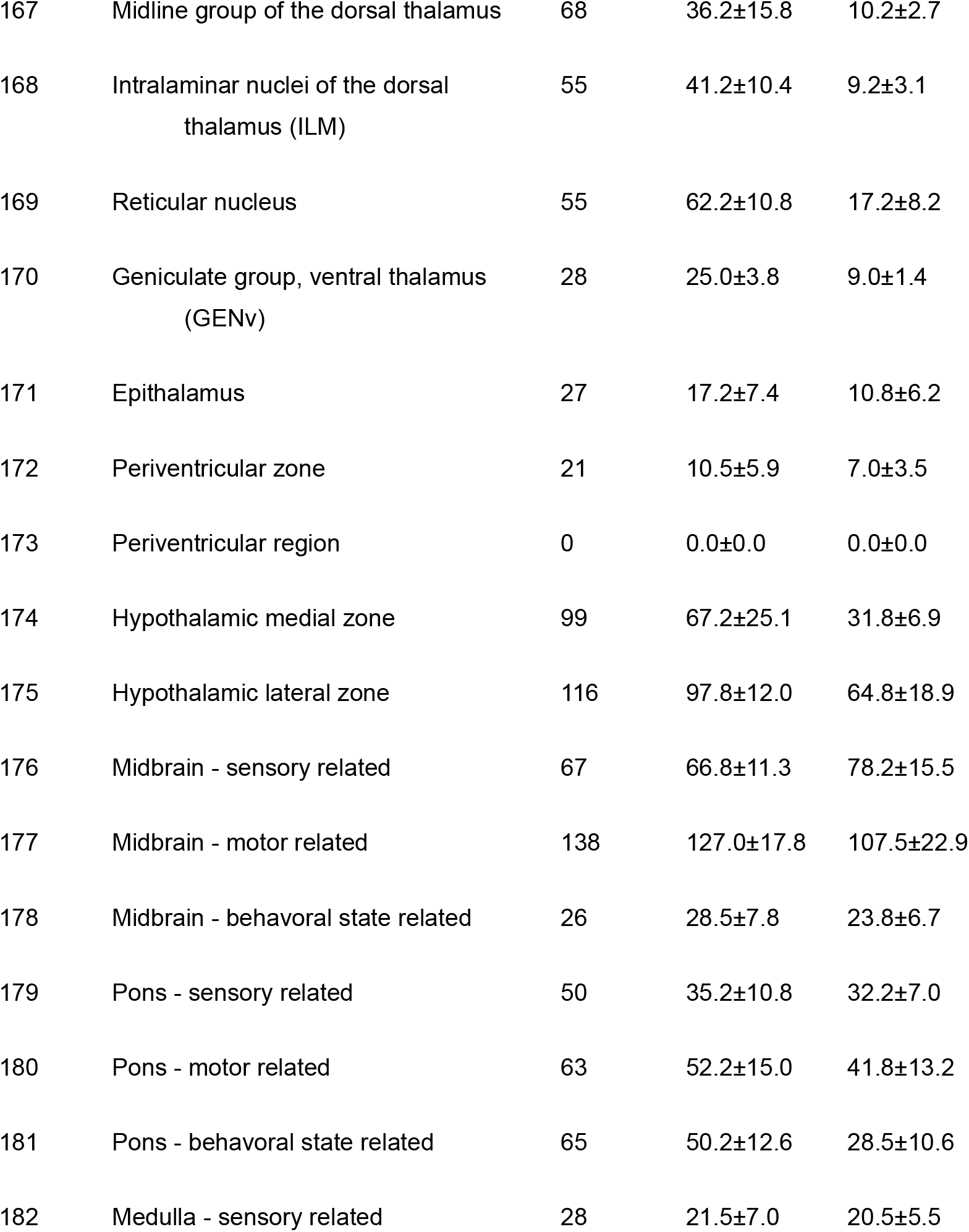
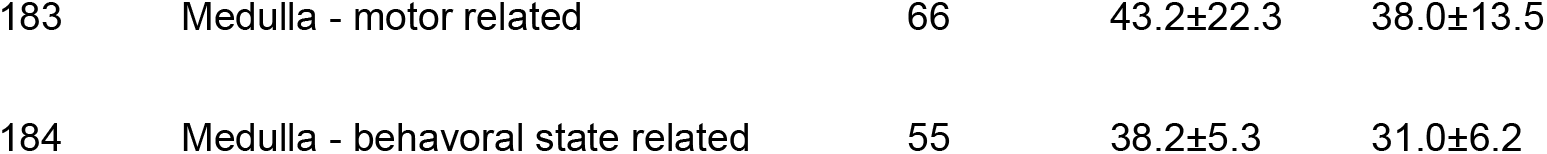
The nodal degrees of each mouse brain region of interest (ROI) in the AMBCA streamline dataset and streamline datasets generated from dMRI data from FiberLM and conventional probabilistic tractography (Prob. Tract.).

